# Mapping the phenotypic repertoire of the cytoplasmic 2-Cys peroxiredoxin – thioredoxin system. 1. Understanding commonalities and differences among cell types

**DOI:** 10.1101/224428

**Authors:** Gianluca Selvaggio, Pedro M. B. M. Coelho, Armindo Salvador

## Abstract

The system (PTTRS) formed by typical 2-Cys peroxiredoxins (Prx), thioredoxin (Trx), Trx reductase (TrxR), and sulfiredoxin (Srx) is central in antioxidant protection and redox signaling in the cytoplasm of eukaryotic cells. Understanding how the PTTRS integrates these functions requires tracing phenotypes to molecular properties, which is non-trivial. Here we analyze this problem based on a model that captures the PTTRS’ conserved features. We have mapped the conditions that generate each distinct response to H_2_O_2_ supply rates (*ν*_sup_), and estimated the parameters for thirteen human cell types and for *Saccharomyces cerevisiae.* The resulting composition-to-phenotype map yielded the following experimentally testable predictions. The PTTRS permits many distinct responses including ultra-sensitivity and hysteresis. However, nearly all tumor cell lines showed a similar response characterized by limited Trx-S^-^ depletion and a substantial but self-limited gradual accumulation of hyperoxidized Prx at high *ν*_sup_. This similarity ensues from strong correlations between the TrxR, Srx and Prx activities over cell lines, which contribute to maintain the Prx-SS reduction capacity in slight excess over the maximal steady state Prx-SS production. In turn, in erythrocytes, hepatocytes and HepG2 cells high *ν*_sup_ depletes Trx-S^-^ and oxidizes Prx mainly to Prx-SS. In all nucleated human cells the Prx-SS reduction capacity defined a threshold separating two different regimes. At sub-threshold *ν*_sup_ cytoplasmic H_2_O_2_ is determined by Prx, nM-range and spatially localized, whereas at supra-threshold *ν*_sup_ it is determined by much less active alternative sinks and μM-range throughout the cytoplasm. The yeast shows a distinct response where the Prx Tsa1 accumulates in sulfenate form at high *ν*_sup_. This is mainly due to an exceptional stability of Tsa1’s sulfenate.

The implications of these findings for thiol redox regulation and cell physiology are discussed. All estimates were thoroughly documented and provided, together with analytical approximations for system properties, as a resource for quantitative redox biology.

**Abbreviations:** ASK1, apoptosis signal-regulating kinase 1; Cat, catalase; GSH, glutathione; GPx1, glutathione peroxidase 1; Grx, glutaredoxin; KEAP1, Kelch-like ECH-associated protein 1; NRF2, nuclear factor erythroid 2-related factor 2; Prx, typical 2-Cys peroxiredoxin; PTTRS, peroxiredoxin / thioredoxin / thioredoxin reductase system; Srx, sulfiredoxin; Trx, thioredoxin; TrxR, thioredoxin reductase.

## Introduction

The PTTRS (Figure 1) plays key roles in antioxidant protection and redox signaling in the cytoplasm of eukaryotic cells. This system controls cytoplasmic hydrogen peroxide (H_2_O_2_) concentrations at low oxidative loads [1–3], and plays prominent signaling roles in vascular adaptation [4], mitogenesis [5], inflammation [6], tumorigenesis [7] and apoptosis [8, 9]. But despite the numerous studies associating the PTTRS to redox signaling, a consensus about the mechanisms conveying H_2_O_2_ signals to redox-regulated targets, and how this system integrates signaling and antioxidant protection is yet to emerge.[10–13] Clarifying how the dynamics of the PTTRS in cells relates to the properties and abundances of these proteins is a critical step towards understanding these problems.

**Figure 1.**
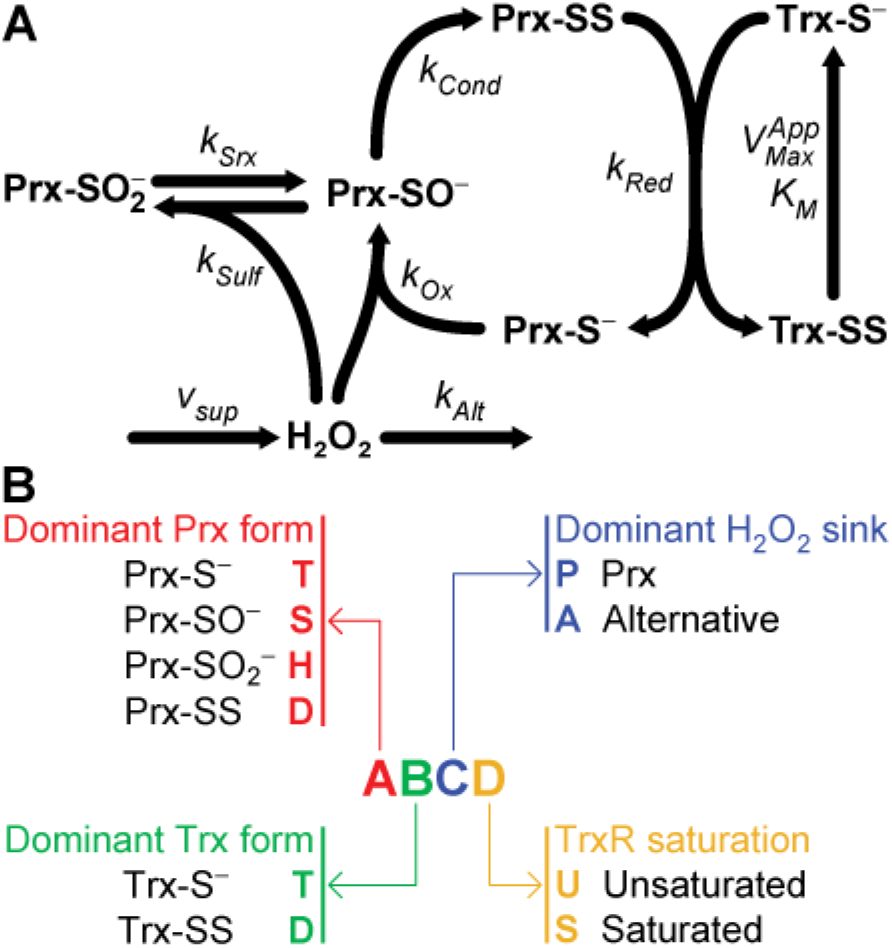
A. A simple model of the peroxiredoxin / thioredoxin / thioredoxin reductase system. The kinetic parameters for each process are indicated near the respective arrow. B. Notation used to designate each phenotypic region.

This dynamics is largely determined by the redox behavior of the cytoplasmic Prx (Figure 1). These are pentamers of dimers in antiparallel orientation. Each monomer carries a very H_2_O_2_-reactive thiolate (peroxidatic cysteine, C_P_) and a less reactive thiol (resolving cysteine, C_R_). H_2_O_2_ oxidizes the peroxidatic cysteine to a sulfenate (Prx-SO^-^), which then condenses with the resolving cysteine of the opposing monomer to form a disulfide (Prx-SS). In eukaryotic Prx the rate of this last step is limited by the local unfolding (LU) around the active site that is required to bring the sulfenate close enough to the resolving cysteine.[12] This delay prompts the accumulation of the sulfenate form, and thereby its further oxidation (called “hyperoxidation”) by additional H_2_O_2_ molecules to sulfinate (Prx-SO_2_^-^) and sulfonate (Prx-SO_3_^-^). The conversion to sulfonate irreversibly inactivates the peroxidatic activity. However, Prx-SO_2_^-^ can be slowly reduced to Prx-SO^-^ at the expense of ATP and reducing equivalents, under catalysis by sulfiredoxin (Srx) [14]. In turn, Prx-SS is reduced by thioredoxin (Trx-S^-^) eventually returning Prx to its fully folded (FF) thiolate form. Thioredoxin becomes oxidized to a disulfide (Trx-SS) in the process, and this disulfide is reduced by NADPH under catalysis by thioredoxin reductase (TrxR).

The characterization of the responses of the PTTRS to H_2_O_2_ in a variety of conditions, cell types and organisms highlighted both commonalities and differences, as the following examples illustrate. Treatment of human erythrocytes (5×10^6^ cells/mL) with H_2_O_2_ boluses up to 200 μM caused both PrxII and Trx1 to accumulate in disulfide form, with virtually no Prx hyperoxidation.[1] Hyperoxidation was detectable only in erythrocytes treated with ≥100 μM H_2_O_2_ boluses after Cat inhibition.[1] In contrast, treatment of Jurkat T cells (10^6^ cells/mL) with ≥100 μM H_2_O_2_ caused extensive hyperoxidation even in absence of Cat inhibition.[1] Remarkably, this happens despite the predominant Prx in Jurkat T cells being PrxI [15], which is more resistant to hyperoxidation than the dominant Prx in erythrocytes, PrxII.[5, 16] Extensive hyperoxidation of PrxI and PrxII was also observed when confluent cultures of human umbilical vein endothelial cells (HUVEC) were exposed to ≥30 μM H_2_O_2_ boluses.[17] In these cells hyperoxidation of PrxII is already extensive, and that of PrxI clearly observable, by 2 min after a 100 μM H_2_O_2_ bolus.[17]

Sobotta *et al.* [18] exposed 10^6^ HEK293 cells/mL to either 0.2 – 3.7 μM H_2_O_2_ steady states for 1 h or 2.5 μM – 5 mM H_2_O_2_ boluses for 5 min. In the former case the fraction of PrxII in disulfide form increased in a dose-dependent manner from ≈5% to 100%. In the latter case that fraction peaked at ≈80% for a 25 μM bolus and progressively decreased for increasing boluses, presumably due to increasing double hyperoxidation of the dimers preventing disulfide formation. Tomalin *et al.* [19] reported that treatment of 2×10^6^ HEK293 cells/mL with 10 – 80 μM H_2_O_2_ boluses for 10 min caused a progressive increase in total hyperoxidation only after a ≈20 μM H_2_O_2_ bolus threshold.

These commonalities and differences prompt important questions with practical relevance for research in thiol redox regulation and for therapy. To what extent are results obtained in one organism or cell type generalizable? What among the many factors that may change form cell type to cell type explain the observed differences in the responses of the PTTRS? What are the determinants of the observed response thresholds? How can oxidative stress and apoptosis be most effectively induced in tumor cells? Will different types of tumor cells react in different ways? Again, understanding the answers to these questions requires clarifying how the dynamics of the PTTRS relates to the properties and cellular abundances of these proteins.

Mathematical modeling has consistently yielded useful insights about the operation of antioxidant and thiol redox systems [20–27] and is recognized as an important tool for the progress of redox biology [28]. Most previous computational studies have focused on accurately modeling specific cells. Instead, the present work seeks to identify generic principles connecting design and function in redox signaling and antioxidant protection by the PTTRS, and on understanding the underpinnings of differences among cell types. Further, we focus on overall dynamic properties, and not yet on details that hinge on further experimental characterization of the components. These distinct goals required a distinct modeling approach. Thus, we proceeded as follows. First, we extensively reviewed the literature and databases to identify (a) the features of the PTTRS in the cytoplasm of eukaryotic cells that are conserved and most relevant for its dynamics, and (b) the typical ranges of the kinetic and composition parameters. This preliminary analysis revealed that most current uncertainties — such as the contribution of GSH for Prx reduction, or a contribution of generic protein thiols for buffering H_2_O_2_ and oxidizing Trx1 — have a minor impact on the overall dynamics of the PTTRS and can be neglected in a first approach. The quantitative analyses supporting this conclusion are documented in the Supplementary Information Section 3 (SI3).

Second, we set up a simple coarse grained model of the PTTRS and analyzed it through a mathematical framework [29, 30] that provides an approximated but intelligible comprehensive description of the relationship between system and molecular properties. The usefulness of this approach to clarify the functional significance of biological variability has been demonstrated.[31] This analysis permitted enumerating the qualitatively distinct states and responses available to the PTTRS and determine closed-form analytical relationships among H_2_O_2_ supply rates, protein concentrations and kinetic parameters that take the system to each state. Importantly, these results do not depend on numerical parameter values, but just on the order-of-magnitude considerations that informed model set up.

Third, based on selected data in the literature and quantitative proteomics datasets we estimated the kinetic parameters and cytoplasmic concentrations for human erythrocytes, hepatocytes, eleven human cell lines, and *S. cerevisiae*. (These estimates are thoroughly documented in the SI.) We validated the quantitative models by comparing computational predictions to the most comprehensive quantitative observations of the PTTRS’ responses to H_2_O_2_ available. In light of the analysis mentioned in the previous paragraph, we then examined the functional implications of the variation in protein composition and properties among cell types, and we dissected the underpinnings of commonalities and differences among the predicted responses.

These analyses show that the PTTRS can *in principle* respond to H_2_O_2_ supply in many distinct ways. Nevertheless, once the actual parameter values and composition are considered distinct tumor cell lines are predicted to show a surprisingly similar response that (a) prevents strong Trx-S^-^ depletion, (b) favors a gradual moderate accumulation of hyperoxidized Prx at high *ν*_sup_, and (c) avoids a run-away hyperoxidation of all the Prx. This response hinges on the Prx-SS reduction capacity just slightly exceeding the maximal steady state Prx-SS production. Its near-universality over cell lines with quite heterogeneous protein composition is due in part to previously undocumented strong correlations between the concentrations of TrxR, Srx and Prx. In turn, erythrocytes, hepatocytes and hepatoma cells are predicted to show a distinct response where at high *ν*_sup_ Trx-S^-^ is depleted and Prx accumulates mainly as Prx-SS.

In all *nucleated* human cells examined the Prx-SS reduction capacity, which is in most cases determined by the TrxR activity, defines a threshold separating two different H_2_O_2_ signaling regimes. At sub-threshold *ν*_sup_ cytoplasmic H_2_O_2_ concentrations are determined by Prx, very low (nM-range) and spatially localized, whereas at supra-threshold *ν*_sup_ H_2_O_2_ concentrations are determined by much less active alternative sinks and in the μM range throughout the cytoplasm. The PTTRS in *S. cerevisiae* is predicted to show a distinct response where at high *ν*_sup_ Prx accumulates in sulfenate form. This is due to the exceptional stability of TSA1’s sulfenate.

The computational predictions are experimentally testable and have important implications for understanding how the PTTRS integrates redox signaling and antioxidant protection, which are examined in the Discussion.

## Model Formulation

We set up a minimal model that captures the basic features of the PTTRS common to most cells where it occurs (Figure 1A). It provides a reasonable description of system behavior in absence of stresses that might deplete NADPH over an extended period. The latter should only occur under strong stresses or in cells with a compromised pentose phosphate pathway. Most healthy cells have glucose 6-phosphate (G6P) dehydrogenase (G6PD) activity in large excess of their capacity to supply G6P, which ensures a fast and effective NADPH supply-demand coupling and avoids strong depletion [32, 33] (SI3.2.9). Indeed, even a 500 μM H_2_O_2_ bolus caused just 30% NADPH depletion to human fibroblasts.[34]

In the model, *ν*_sup_ stands for the overall supply of H_2_O_2_ to the cytoplasm. H_2_O_2_ can be reduced by Prx through both Prx-S^-^ (rate constant *k*_Ox_) and Prx-SO^-^ (*k*_Sulf_). The rate constant *k*_Ox_ can be regarded as an *effective* rate constant for H_2_O_2_ reduction by Prx-S^-^, which allows to consider a diminished effective peroxidatic activity such as postulated [21] for PrxII in human erythrocytes. The H_2_O_2_ can also be cleared by efflux from the cytoplasm, by the activities of catalase, peroxidases, 1-Cys Prx, and by reaction with other thiolates. These alternative sinks were aggregated into a single process with first-order kinetics (rate constant *k*_Alt_), as peroxidases should only saturate at very high (>17 μM) intracellular H_2_O_2_ concentrations (SI3.2.2.1). With the exception of erythrocytes [21], under low *ν*_sup_ these alternative sinks contribute little for H_2_O_2_ clearance in all other cells analyzed (SI3.2.2, Supplementary Table, ST, 6).

For simplicity, we consider each Prx’s proximate C_P_-C_R_ pair as an independent functional unit. The Tpx1 peroxiredoxin of *Schizosaccharomyces pombe* [19] and the human mitochondrial Prx3 [35] may violate this assumption. However, there is no evidence that the same happens to the same extent for other eukaryotic Prx in the cytoplasm [35]. Further, even in the noted exceptions, the PTTRS shows qualitatively the same overall dynamics.

The FF → LU transition plus condensation sequence, as well as the Prx-SS reduction plus LU → FF transition were each aggregated into single steps, with apparent first- and second-order rate constants *k*_Cond_ and *k*_Red_, respectively.

The reduction of Prx-SO_2_^-^ to Prx-SO^-^ is treated as a pseudo-first-order process (*k*_Srx_). Srx has relatively low Michaelis constants for ATP and Trx1-S^-^ [36], which supports this approximation. Its Michaelis constant for Prx-SO_2_^-^ has not been determined. However, from results in ref. [37] it can be inferred that the *K_M_* of *Saccharomyces cerevisiae* Srx for Tsa1 is ≈20 μM. This suggests that Prx-SO_2_^-^ reduction by Srx follows pseudo-first order kinetics with respect to this substrate *in vivo,* except under high oxidative stress.

The reduction of Trx-SS was treated as a one-substrate Michaelis-Menten process. This process is catalyzed by TrxR and follows a ping-pong mechanism that uses NADPH as a second substrate [38]. However, the *K*_MTrxR,NADPH_ of human TrxR1 is 6 μM [39] and the apparent *K*_MTrXR,NADPH_ is substantially lower when the enzyme is far from saturation with Trx-SS.

Therefore, physiological concentrations of NADPH should be saturating for TrxR, which justifies neglecting the influence of this substrate on the reaction rate. Below we write the rate expression for TrxR as function of a 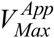 to highlight that the maximal rate of TrxR may decrease at very high *ν*_sup_ due to NADPH depletion or to TrxR inactivation by electrophilic lipid peroxidation products [40].

PrxI can be glutathionylated at C52, C173 and C83, and the extent of glutathionylation was 40%-60% increased (vs. untreated controls) 10 min after treatment of A549 or HeLa cells with a 10 μMH_2_O_2_ bolus.[41] Both Srx and glutaredoxin 1 (Grx1) can catalyze the deglutathionylation of these residues.[41] Altogether, these findings suggest that glutathione (GSH) might reduce PrxI-SS and/or PrxI-SO^-^. This hypothesis was recently supported by the observation that GSH plus Grx1 can reduce both PrxII-SS and PrxII-SO^-^.[42] However, as discussed in SI3.2.4 these reactions do not dominate the dynamics of the PTTRS and can thus be neglected in a coarse-grained model for cells where PrxI is the dominant cytoplasmic Prx. They can play a significant role in increasing resistance to hyperoxidation and reducing Prx-SS in erythrocytes, where PrxII is the dominant Prx and the TrxR activity is very low.[42] But even here only a very minor fraction of PrxII is glutathionylated.[42]

Trx-SS can also be reduced by GSH + Grx1, but similarly to the case for Prx this is not the dominant reductive process [43], and can thus be neglected in a coarse-grained model of the PTTRS.

Although Trx1-S^-^ can be oxidized by numerous protein disulfides and several enzyme-catalyzed processes, our estimates (SI3.2.7) indicate that under oxidative stress Prx-SS reduction is the dominant process oxidizing Trx1-S^-^. In turn, at lower *ν*_sup_ cells’ TrxR activity is sufficient to keep

Trx1 strongly reduced despite all the oxidizing processes. Because the objective for the model is to provide a simple description of the PTTRS dynamics and not a quantitative estimation of the Trx redox status we neglected the other processes oxidizing Trx-S^-^. For similar reasons we also neglected the oxidation of Prx-S^-^ by Trx-SS.

Note that the overall reaction for the chemical system represented in this model couples the reduction of H_2_O_2_ to water to NADPH oxidation. The overall process is extremely thermodynamically favorable owing largely to the high redox potential of the first half reaction [+1.32 V at 25 ^o^C, pH 7 [44]]. Therefore, the PTTRS should operate irreversibly.

The assumptions discussed above and the scheme in Figure 1A translate into the following system of algebraic differential equations to describe the dynamics of the PTTRS:

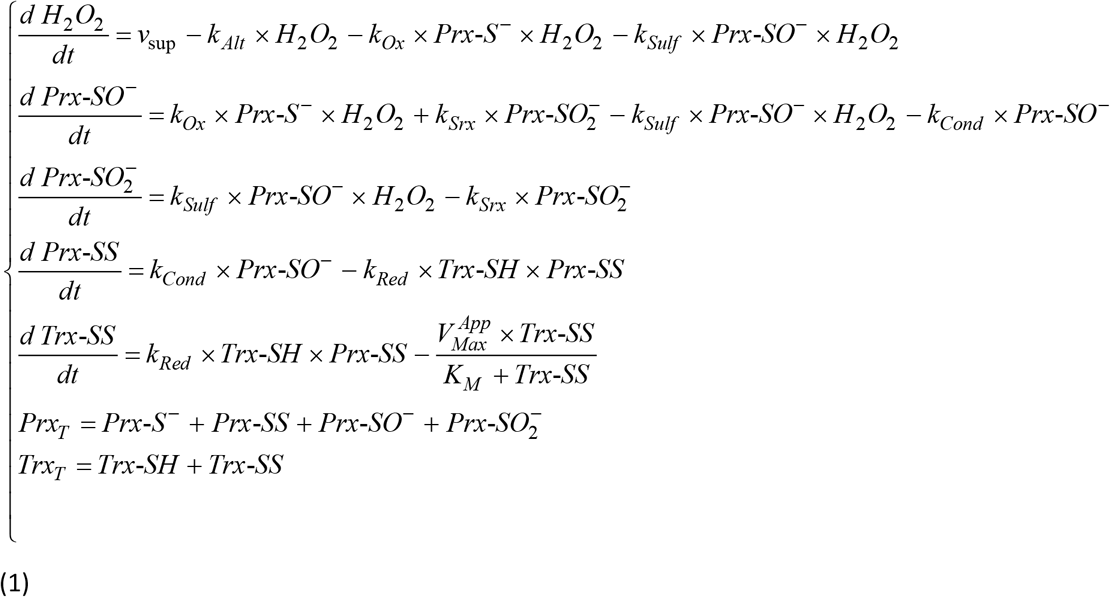

Henceforth, we will denote this model by “Model 1”.

For physiological Prx concentrations there are substantial concentration gradients of H_2_O_2_, Prx-SO^-^, and Prx-SS over the cytoplasm [13, 22, 45], which this model does not account for. However, recent reaction-diffusion simulations [13] confirm that the model provides a good overall description of the behavior of the average concentrations of these species.

## Methods

The systems design space analysis was performed using the algorithms described in SI1, which extends the applicability of the approach published in ref. [30]. Sensitivity analyses were performed according to the methodology in refs. [46, 47]. Except where otherwise stated, all these analyses and the numerical simulations were performed in *Mathematica*^TM^ 11 [48].

Parameters for human erythrocytes were estimated as described in ref. [21]. Those for *Saccharomyces cerevisiae* and for human Jurkat T, A549, GAMG, HEK293, HeLa, HepG2, K562, LnCap, MCF-7, RKO, and U-2 OS and hepatocyte cells were estimated from the literature and databases as described in SI3.

Numerical simulations for a quantitative comparison between computational and experimental results on PrxII were based in the two-Prx model described in SI4.

## Results

### A phenotypic map of the PTTRS

We first seek to map the properties of the system as a function of kinetic parameters and protein concentrations. As a starting point, this requires analyzing the steady state solutions of Model 1. However, these solutions cannot be expressed in closed analytical form, and the large number of parameters prevents an effective numerical exploration. We therefore applied the system design space methodology [29, 30] to obtain an intelligible approximate description. This methodology subdivides the parameters space into a set of regions. The dynamics in each region is described by a distinct combination of alternatively dominant production and consumption fluxes for each chemical species, and of alternatively dominant concentrations among the forms included in each moiety-conservation cycle. Whenever a region contains a steady state solution, this is guaranteed to be unique and analytically described by a simple product of power laws of the parameters. By the construction of the approximation, these regions represent qualitatively distinct behaviors of the system, and are accordingly denoted by *“phenotypic regions”.* The parameters space partitioned into the set of phenotypic regions is called the system’s *“design space”.*

The construction of the design space for Model 1 is explained in SI1,2. This design space contains 13 regions with positive steady state solutions, but not all of these are biologically plausible. In order to select these, one has to consider the ranges of kinetic parameters and protein concentrations found in real cells. We consider the following three plausibility criteria cumulatively (see SI2 for mathematical definitions).

First, the maximum flux of Prx-SO_2_^-^ reduction is the lowest maximum flux of the system. This is because Srx is an inefficient enzyme [37,49–51] and is much less abundant in cells than the other proteins considered in the model (ST6).

Second, the pseudo-first-order rate constant for H_2_O_2_ reduction by Prx-S^-^ exceeds the rate constant for Prx-SO^-^ condensation. This follows from the high reactivity and abundance of typical 2-Cys Prx in the cytoplasm, contrasting with the kinetic limitation imposed by the local unfolding to condensation step.

Third, Prx sulfinylation is the slowest among all (aggregated) H_2_O_2_-consuming processes in Model 1. This holds with a large margin for all cells we examined (SI3).

Only the 8 phenotypic regions that we describe below satisfy the three plausibility criteria above. We designate each region by a four-character code as explained in Figure 1B. A pictorial representation of system steady state properties in each of these regions is presented in Table 1. For the corresponding mathematical descriptions of the steady state properties and of the boundaries of each region in the parameters space please see ST1 and ST2, respectively, in SI2.1.

**Table 1.**
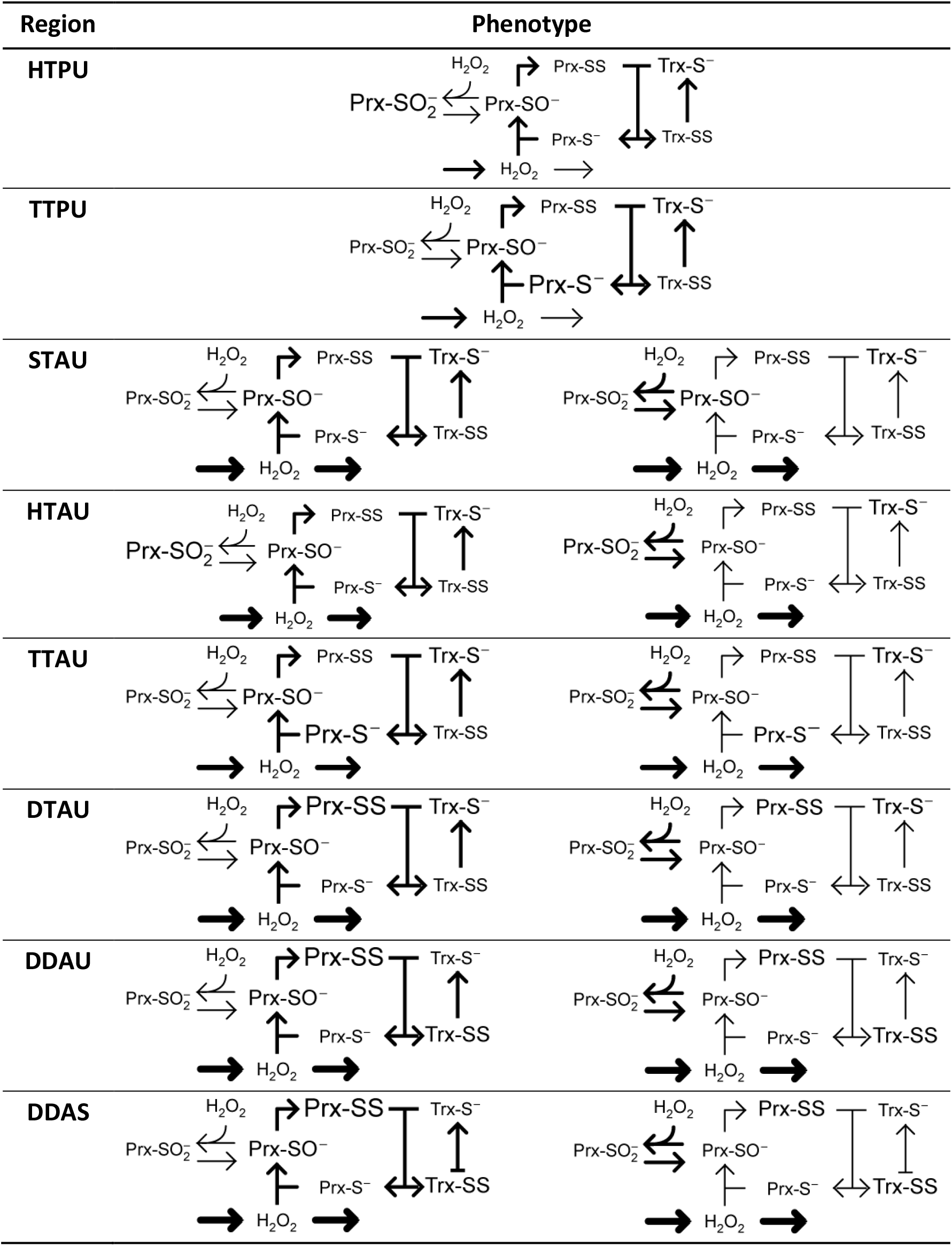
System states in the physiologically plausible phenotypic regions. Each phenotypic region is characterized by a hierarchy of concentrations of alternative protein forms (represented by symbols of different sizes) and one or more flux hierarchies (arrows of different widths). Distinct flux hierarchies belong to the same phenotypic region when they originate the same steady state solution. A bar at a reactant tip of an arrow indicates that the reactant in point is saturating for the corresponding process.

Phenotypic regions TTPU and TTAU are characterized by the thiol(ate) forms of Prx and Trx being the dominant ones and differ on whether most of the H_2_O_2_ is consumed by Prx (TTPU) or by alternative sinks (TTAU) (Table 1). These regions occur where, cumulatively, the H_2_O_2_ supply is low and the TrxR activity is not too low (ST2, Figure 2). In these regions, the concentrations of Prx-SO^-^, Prx-SS and Trx-SS show an approximately linear response to changes in *ν*_sup_ (ST1, Figure 2).

**Figure 2.**
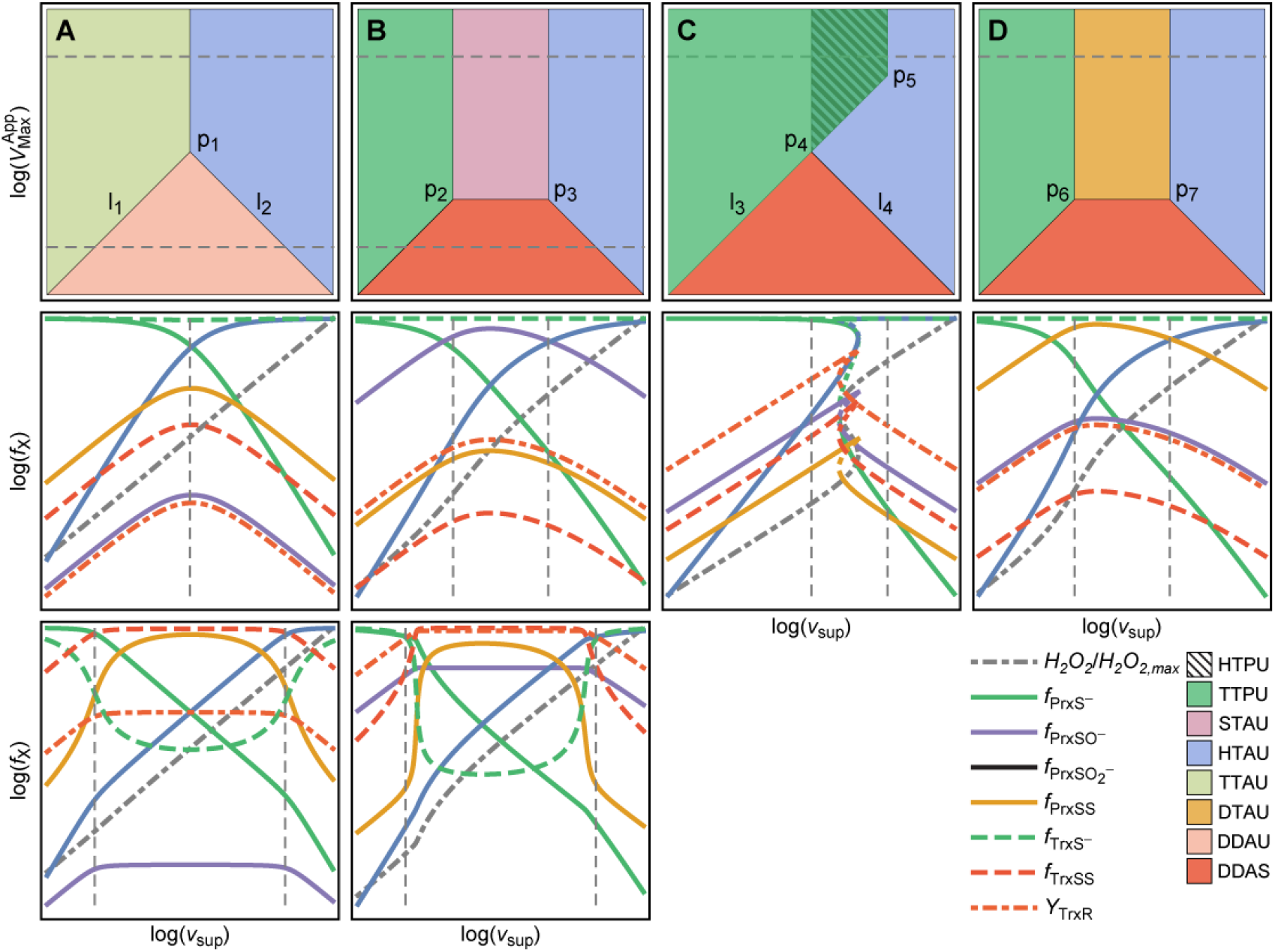
Slices of the design space of the PTTRS over (*ν*_sup_,V_Max_^App^) planes, showing the relative locations of the various phenotypic regions for distinct cell compositions. The second and third rows show the responses of the fractions of Prx and Trx in each form, TrxR saturation and cytoplasmic H_2_O_2_ concentration to *ν*_sup_ for the V_Max_ values marked by the dashed horizontal lines in panels A-D. The vertical dotted lines mark region boundaries. H_2_O_2_ concentration is scaled by maximal value achieved in each plot. Each region is characterized by distinct concentration hierarchies as well as by distinct dependencies on H_2_O_2_ supply. Boundaries among phenotypic regions correspond approximately to crossover points where these concentration hierarchies or TrxR saturation qualitatively change. The composition ranges yielding each type of section are as follows:

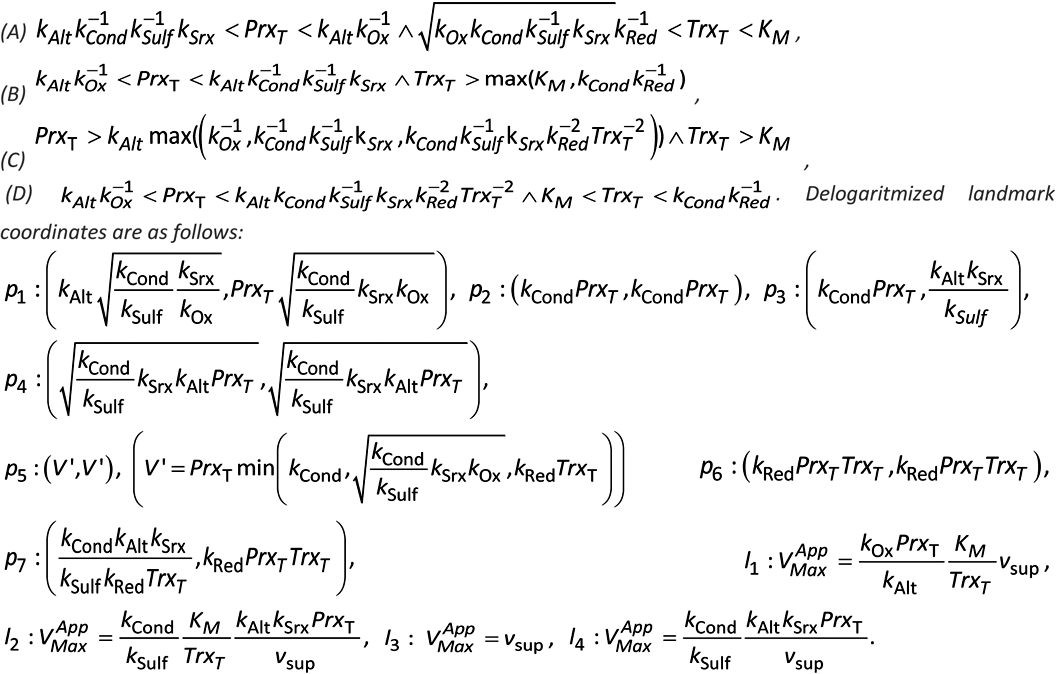 For multistationarity to occur the ratio between the highest and the lowest value of ν_sup_ in the overlap region in (C) must exceed 2.

Region HTAU is characterized by extensive Prx sulfinylation and low Trx oxidation (ST1, Figure 2). This region occurs at very high *ν*_sup_ and not very low TrxR activity (ST2, Figure 2).

Under some conditions, regions TTPU and HTAU partly overlap (Figure 2C). When this occurs, there is also a region (HTPU) of unstable steady states that coincides with the overlap between TTPU and HTAU. This feature reveals the possibility of bi-stability and hysteresis in the PTTRS, which had not been appreciated before. That is, under these conditions as *ν*_sup_ increases up to the critical value 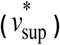 at the right hand side of region TTPU, entering region HTAU, the concentrations of Prx-SO_2_^-^ and H_2_O_2_ abruptly increase, whereas those of Prx-S^-^, Prx-SO^-^, Prx-SS, and Trx-SS abruptly decrease. However, as *ν*_sup_ decreases from high values in region HTAU the opposite transition occurs only at a lower critical value 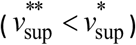 at the left hand side of region HTAU, entering region TTPU. The abrupt changes are driven by the following positive feedback. As *ν*_sup_, and hence H_2_O_2_ concentration, increases Prx becomes more hyperoxidized, depleting Prx-S^-^. As reduction by Prx-S^-^ is the main H_2_O_2_ sink, this depletion causes a sharp increase in the concentration of H_2_O_2_, which accelerates hyperoxidation even further. Biological systems use toggle switches like this to avoid random back-and-forth switching between discrete physiological or developmental states, driven by gene expression noise and environmental fluctuations.[52] On the other hand, the positive feedback just described will cause near-complete Prx hyperoxidation, from which cells can recover only very slowly due to the usually very low Srx activity. Therefore, the findings above raise the intriguing question of whether cells use the PTTRS’ hysteretic behavior as part of a stress switch, or avoid it due to its potentially very deleterious consequences.

Regions STAU and DTAU are characterized by Trx being predominantly in thiol form and the dominant Prx forms being Prx-SO^-^ or Prx-SS, respectively (Table 1, Figure 2B,D, ST1). Both regions occur at intermediate H_2_O_2_ supplies and high TrxR activities, only under conditions where regions TTPU and HTAU do not overlap (ST2, Figure 2B,D).

Finally, regions DDAU and DDAS are characterized by the dominance of the disulfide forms of both Prx and Trx, and differ on whether TrxR is saturated (DDAS) or not (DDAU) (Table 1, ST1). They occur at intermediate H_2_O_2_ supplies and low TrxR activities (Table 1, Figure 2, ST2).

The analysis of the design space permits the following generalizations. First, the system can always be driven to either TTPU or TTAU by making *ν*_sup_ sufficiently low, and to HTAU by making *ν*_sup_ sufficiently high. However, the latter high *ν*_sup_ values are not necessarily physiological. Second, the system can always be driven to regions DDAU and DDAS through a strong enough inhibition/under-expression of TrxR or Trx, though DDAS becomes unreachable where the concentration of Trx is too low to saturate TrxR. Third, not all dominance configurations are feasible or biologically plausible. For instance, whenever Trx-SS is the dominant Trx form Prx-SS is the dominant Prx form, and whenever Prx-SO_2_^-^ is the dominant Prx form Trx-S^-^ is the dominant Trx form. Fourth, only regions TTPU, HTPU and HTAU can overlap. In particular, regions TTAU and HTAU cannot overlap, which means that the hysteretic behavior described above can only occur when Prx are the main H_2_O_2_ consumers under low oxidative loads.

### The PTTRS permits many distinct responses to H_2_O_2_ supply

As illustrated in Figure 2 the eight phenotypic regions can be arranged in several alternative ways, yielding potentially many qualitatively distinct responses of the various chemical species to changes in *ν*_sup_ and 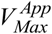. We have systematically inventoried the qualitatively different arrangements of the eight phenotypic regions in the 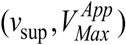 plane that are possible under physiologically plausible conditions (SI2.2). This analysis reveals that 12 qualitatively distinct generic configurations can occur (SF2). Furthermore, 10 qualitatively distinct generic types of responses to *ν*_sup_ are possible, corresponding to as many different sequences of phenotypic regions. (“Generic” here means that it does not hinge on specific “pointwise” relationships between parameters.) Six of these generic types of responses are illustrated in Figure 2, and the conditions where each of the 10 responses hold are listed in ST3.

With relevance for signal transduction, these results demonstrate that the PTTRS can exhibit many distinct types of responses to *ν*_sup_. These include: (i) saturation, as for *Trx-SS* and *Prx-SO^-^* transitioning from TTAU to DDAU (Figure 2A); (ii) ultrasensitivity, as for *Prx-SS, Trx-SS* and *Trx-S^-^* transitioning from TTPU to DDAS and from DDAS to HTAU (Figure 2B); (iii) hysteresis (toggle-switch behavior), as for all variables transitioning from DTAU to HTAU over the overlap region (Figure 2C); (iv) non-monotonic behavior of all variables except *H_2_O_2_*, *Prx-S*^-^ and *Prx-SO*_2_^-^.

### Composition and kinetic parameters of the PTTRS show high variability across cell types

All the results obtained above depend on just a few generic assumptions about parameter values and protein concentrations. They suggest a potentially high diversity of responses of the PTTRS to H_2_O_2_ supply under the biological plausibility conditions specified above. But do cells manifest all this diversity, or do they orchestrate protein concentrations and kinetic parameters to generate a common phenotype? That is the problem we address over the next sections.

In order to address the previous question and to quantitatively define region boundaries and their locations relative to physiological operating ranges, parameter values must be known. Until recently, this requirement would pose an insurmountable obstacle. However, a consistent effort by several laboratories has provided reliable determinations of most of the necessary kinetic parameters, and quantitative proteomic methodologies now allow reliable estimates of the concentrations of all the relevant PTTRS proteins. These developments permitted estimating the relevant parameters for human erythrocytes (as per ref. [21]) and hepatocytes, one non-cancer human cell line (HEK293), ten human tumor cell lines, and *S. cerevisiae.* Protein abundances for the eleven human cell lines were determined in a single laboratory through the same state-of-the-art methods [53], which adds confidence on the comparability of the results. Furthermore, a separate quantitative proteomics study [54] obtained protein abundances for human hepatocytes and for the HepG2 cell line under the same conditions. The estimates are documented in SI3, and parameters for all cell types are presented in Table 2. A more detailed account of the estimated protein concentrations and activities contributing for these aggregated parameters is presented in ST5. The latter results suggest that at low oxidative loads the 2-Cys peroxiredoxins consume ~99% of the H_2_O_2_ in the cytoplasm of human cells. In turn, glutathione peroxidase 1 (GPx1) is much less abundant and typically consumes <0.2% of the H_2_O_2_, and Cat has a negligible contribution because it is located in peroxisomes. The main contributor for cytoplasmic H_2_O_2_ clearance other than the typical 2-Cys peroxiredoxins is PrxVI. Despite their very high PrxII concentrations, erythrocytes are an exception to this pattern, as experimental evidence [2,55–57] indicates that Cat plays a major role in H_2_O_2_ clearance in these cells even at modest H_2_O_2_ supply rates. This is due to a very high cytoplasmic concentration of Cat and presumably to a postulated [21] strong and quickly reversible PrxII inhibition Cat.

**Table 2.**
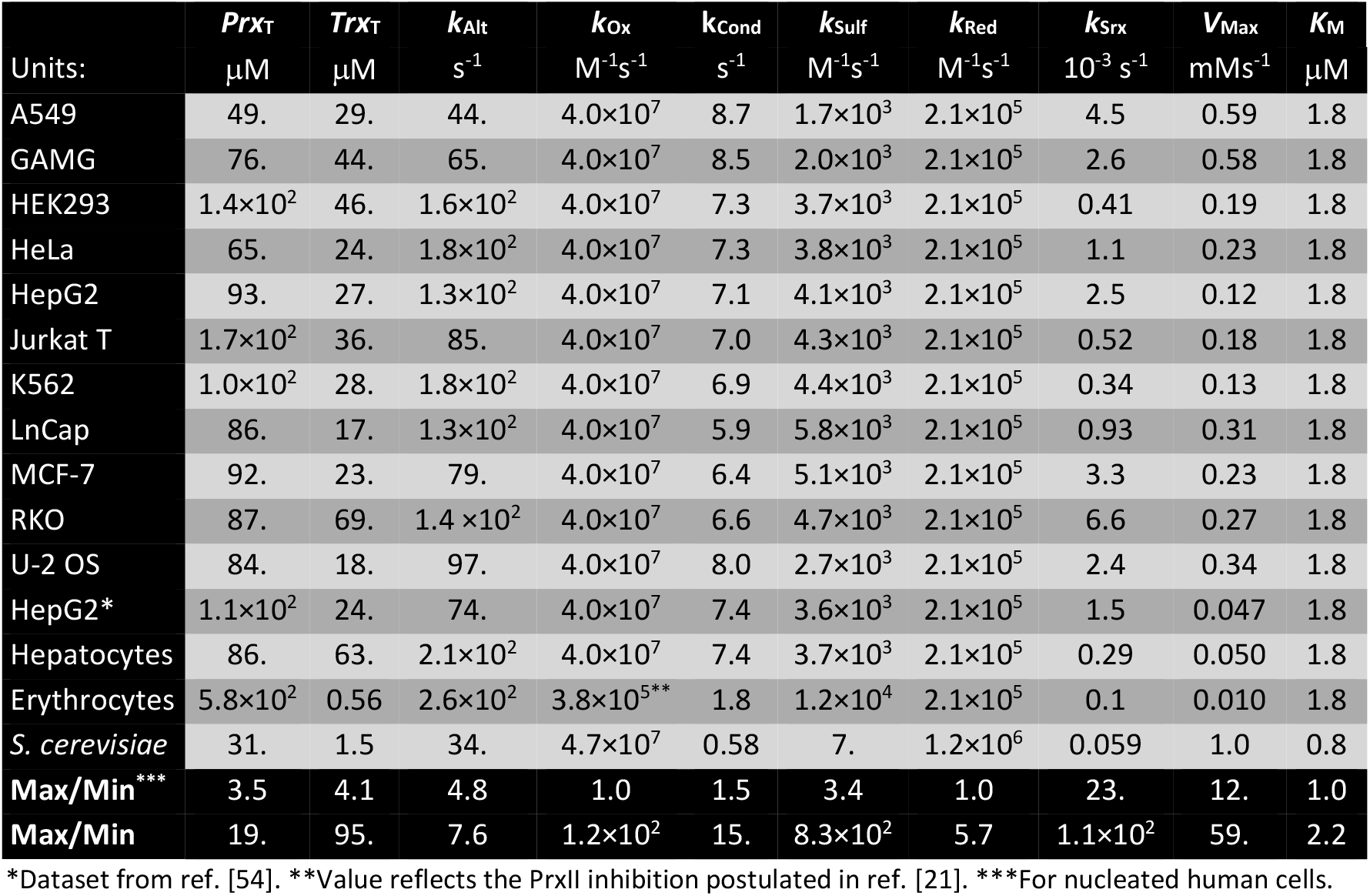
Summary of estimated parameters for the cell types addressed in this work.

The last two rows in Table 2 highlight that protein concentrations and kinetic parameters of the PTTRS can vary by several orders of magnitude over the various cell types and organisms. Remarkably, both Srx and TrxR activities can vary by over one order of magnitude among the nucleated human cells in our dataset. This variability makes the question about phenotypic diversity especially pertinent.

### The underlying model is consistent with and yields insight on experimental observations

Before analyzing the predicted phenotypes for the various cells one must test if the underlying model and parameter estimates yield predictions that are consistent with known experimental observations. Few studies so far have investigated the responses of the PTTRS quantitatively under steady state conditions. However, experimental observations of cells’ responses to H_2_O_2_ boluses can also be used to validate model and estimates.

Sobotta *et al.* [18] examined the response of PrxII redox status of HEK293 cells exposed to 0.2 – 3.7 μM H_2_O_2_ steady states and to 5 min 2.5 – 5000 μM H_2_O_2_ boluses. Because PrxI is more abundant than PrxII in these cells (ST6) the *Prx-SS* variable in Model 1 reflects mainly the concentration of PrxI-SS. Seeking a more direct quantitative comparison between computational predictions and these experimental observations we set up the model presented in SI4. This model (“Model 2”) differs from Model 1 by considering the redox cycles of PrxI and PrxII separately and by considering H_2_O_2_ exchange between the medium and the cytoplasm explicitly. Parameterized with the data from Table 2 and ST6 for HEK293 cells, Model 2 shows qualitative agreement with the experimental observations (SF9). Namely, a near-linear increase in PrxII-SS with extracellular H_2_O_2_ concentration up to near-maximal values and the biphasic response of PrxII-SS to H_2_O_2_ boluses.

However, the maximal fraction of PrxII disulfide attained is much lower than observed. The design space analysis above helps tracing the likely source of this discrepancy. The Prx-SS response curves in Figure 2 show that for high fractions of Prx crosslinking to be achieved with a near-linear response to *ν*_sup_ as observed, the protein composition should allow attainment of region DTAU (*e.g.* Figure 2D). The analysis shows that this will happen if the total concentration of Trx is lower than assumed. Indeed, numerical simulations of Model 2 with the cytoplasmic concentration of Trx1 set to ≈1.5 μM (≈3% of the previously estimated value) show remarkable agreement with the observations (SF10A).

Model 2 with *Trx_T_* = 1.5 μM also agrees with the experimental observations [19] of a threshold H_2_O_2_ bolus beyond which Prx hyperoxidation starts increasing, and intracellular H_2_O_2_ increases more steeply (SF10B,C). This threshold behavior has been attributed to the saturation of alternative H_2_O_2_ sinks and a buffering effect by generic protein thiols.[19] However, it is predicted even by Model 1, which includes neither of these effects. A comparison of the simulated progress curves of the various species for boluses around the threshold value (SF11) points to the following alternative explanation. The steeper increase in hyperoxidation starts once the bolus becomes sufficient to fully oxidize Prx to Prx-SS, thereby sharply decreasing the cytoplasmic H_2_O_2_ clearance rate. In turn, the ensuing sharp increase in cytoplasmic H_2_O_2_ concentration causes a strong increase in sulfinylation and accumulation of Prx-SO_2_^-^. Noting that the full oxidation of the Prx-S^-^ pool to Prx-SS can only happen when the rate of Prx-S^-^ oxidation exceeds the rate of Prx-SS reduction, one can make several experimentally testable predictions about the factors that influence the threshold value of extracellular H_2_O_2_. First, the threshold must be inversely proportional to the membrane permeability (SF12A,B), as for the same extracellular H_2_O_2_ the H_2_O_2_ influx rate is proportional to membrane permeability. There is growing evidence that cells’ permeability to H_2_O_2_ is largely determined by some aquaporins [58, 59], and the previous prediction can thus be tested through the use of aquaporin inhibitors. Second, the threshold should be higher the higher the concentration of Trx available in the cytoplasm (SF12C,D), as Prx-SS reduction in these cells is rate limited by Trx availability. Third, increasing the TrxR activity should have qualitatively the same effect as increasing the Trx concentration, but the effect should be much less pronounced in these cells than the latter (SF12C,D) because here the TrxR activity is not rate-limiting for Trx-SS reduction. In most other cells examined TrxR activity is rate-limiting for Trx-SS reduction in absence of extensive Trx1 sequestration and should there have a stronger effect on the threshold. Fourth, the activity of alternative H_2_O_2_ sinks should have little effect on the threshold in these cells (SF12E,F). This because while there is Prx-S^-^ available those sinks contribute little for H_2_O_2_ elimination and to determine the cytoplasmic H_2_O_2_ concentration. Fifth, the total concentration of Prx should have virtually no effect on the threshold (SF12E,F). This because as Prx-S^-^ consumes the overwhelming majority of the H_2_O_2_, the rate of Prx-SS formation is determined by the rate of H_2_O_2_ supply and not influenced by *PrX_T_*. The threshold’s dependence on the last four factors was not tested for HEK293 cells. However, the observations reported in ref. [19] for *Schizosaccharomyces pombe* agree with all these predictions, which further supports our interpretation of the threshold’s underpinnings.

Further experimental data for model validation can be found in Low *et al.* [1], where the response of PrxII redox status in Jurkat T cells and erythrocytes to various H_2_O_2_ boluses is examined. Model 2, parameterized with the data from Table 2 and ST6 for Jurkat T cells yields results in very good agreement with the experimental observations for these cells (SF14). A much sharper threshold response of PrxII-SO_2_^-^ than for HEK293 cells is predicted, which the experimental observations do indeed support. Remarkably, in the case of Jurkat T cells setting the Trx concentration to 3% or even 30% of the value estimated from the quantitative proteomics data yields a poor agreement with the experimental observations, which indicates that Trx1 is not much sequestered in these cells.

Because in erythrocytes PrxII is by far the dominant Prx, one can use Model 1 to simulate the redox state of PrxII. Parameterizing Model 1 with the data from Table 2 and ST6 for these cells again yields results in very good agreement with the experimental observations (SF15).

Although some of the simulations above were carried out with Model 2, the single-Prx Model 1 provides a good approximation of the overall Prx redox state (SF16). Thus, altogether, the results above validate the model and parameter estimates underlying the subsequent analysis, though the possibility that in some cells just a small fraction of the Trx1 is available to reduce Prx-SS needs to be considered. To obtain a more general perspective of the predicted phenotypes of the cell types under consideration we now turn again to the design space analysis, now informed by the quantitative estimates.

### Most human cell lines share a common PTTRS phenotype

When all the Trx1 is considered available to reduce Prx-SS the slices of the quantitative design spaces in the plane 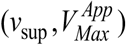 (Figure 3, SF17), and the responses to *ν*_sup_ (Figure 4, SF18) for all cell types reveal the following remarkable patterns.

**Figure 3.**
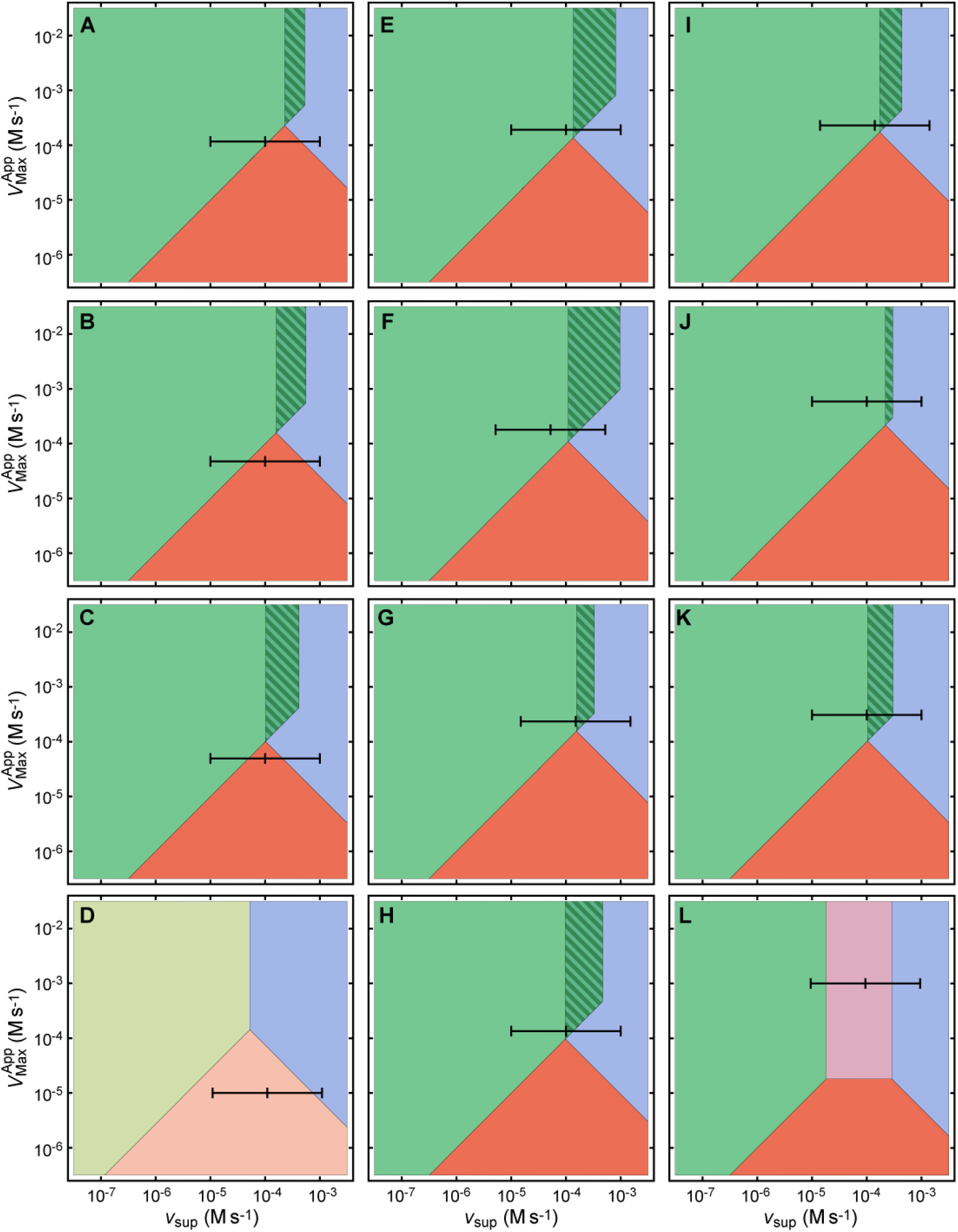
Slices of the design space of the PTTRS over the physiological (ν_sup_,V_Max_^App^) plane for human cell types and S. cerevisiae as computed for the parameters estimated in ref. [21] and SI3. The black scales inside the plots mark the apparent V_Max_ of TrxR and the values of ν_sup_ corresponding to 1 μM, 10 μM and 100 μM extracellular H_2_O_2_. These values of ν_sup_ were estimated based on the known cell permeability and morphology (HeLa, MCF-7, Jurkat T cells, erythrocytes and S. cerevisiae) or assuming kı_n_f = 10 s^-1^ (all other cells). Note the logarithmic scales. Color codes are as for Figure 2. A, HepG2, ref. [53]; B, HepG2, ref. [54]; C, hepatocytes; D, erythrocytes; E, HEK293; F, Jurkat T; G, HeLa; H, K562; I, MCF-7; J, A549; K, LnCap; L, S. cerevisiae. Design space slices for GaMG, RKO and U-2 OS cells are presented in SF17.

**Figure 4.**
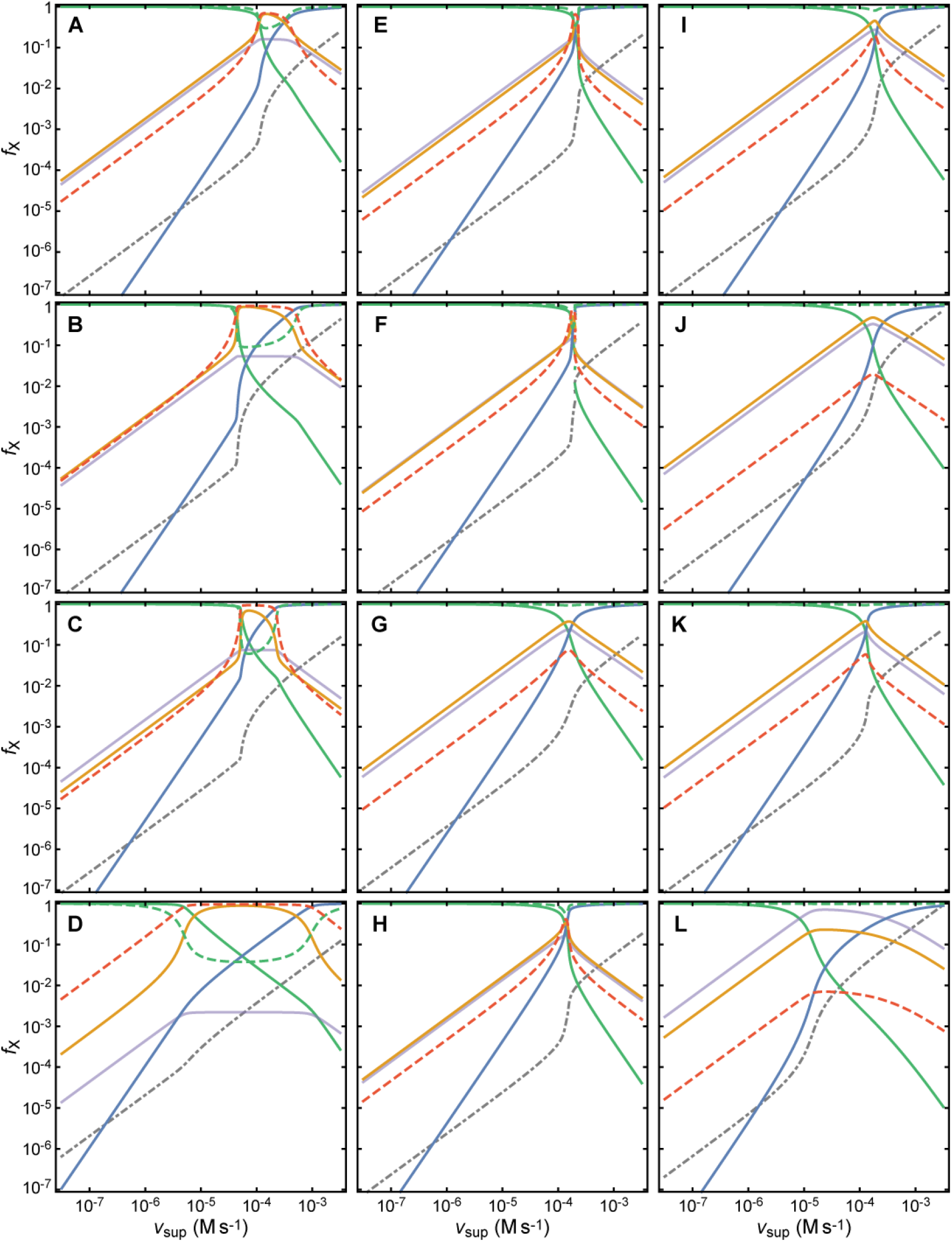
Responses of the PTTRS to H_2_O_2_ supply rates for human cell types and S. cerevisiae as computed for the parameters estimated in ref. [21] and SI3. Note the logarithmic scales. The plots were obtained by numerical integration of equations (1) with the parameters in Table 2. Predictions of the responses at ν_sup_ > ⅛0.5 mM s^-1^ may be inaccurate due to neglect of NADPH depletion. Color codes are as for Figure 2, except that cytoplasmic H_2_O_2_ concentrations are scaled by 100 μM. A, HepG2, ref. [53]; B, HepG2, ref. [54]; C, hepatocytes; D, erythrocytes; E, HEK293; F, Jurkat T; G, HeLa; H, K562; I,MCF-7; J, A549; K, LnCap; L, S. cerevisiae. Responses of GaMG, RKO and U-2 OS cells are presented in SF18.

First, the design spaces based on two quantitative proteomic datasets obtained in distinct laboratories [53, 54] for the HepG2 cell line are remarkably consistent (Figure 3A,B). Namely, they both predict that at relatively wide range of intermediate values of *ν*_sup_ (≈40 – 450 μM s^-1^) both Prx and Trx accumulate mainly in disulfide form (Figure 4A,B). This accumulation sets in at a sharp *ν*_sup_ threshold and is accompanied by a sharp ultrasensitive increase in the H_2_O_2_ concentration. Prx-SO_2_^-^ only becomes the predominant Prx form at even higher *ν*_sup_. This behavior matches response PDS in ST3.

Second, the design space (Figure 3C) and response to *ν*_sup_ (Figure 4C) for hepatocytes are very similar to those for the hepatoma-derived HepG2 cell line. Thus, the distinctive compositional pattern of hepatocytes (Table 2, ST6) does not translate into qualitative differences from HepG2 cells in the response of the PTTRS to H_2_O_2_.

Third, the responses of human erythrocytes to H_2_O_2_ supply (Figure 3D, Figure 4D) are predicted to show some unique features, characteristic of Response A in ST3. Namely, at low *ν*_sup_ the PTTRS of erythrocytes operates in phenotypic region TTAU, and not in TTPU as all the other cells examined. Also unlike all the other cells, in erythrocytes the concentration of Trx1 is insufficient to saturate TrxR, and therefore at higher *ν*_sup_ the PTTRS operates in region DDAU and not in region DDAS. Consequently, the Trx-SS concentration should not show the sharp ultrasensitive behavior predicted for hepatocytes and HepG2 cells, and the H_2_O_2_ and Prx-S^-^ concentrations also should not show the strongly ultrasensitive behavior predicted for all other human cells in this study. The response of the PTTRS in erythrocytes is otherwise most similar to that of hepatocytes and HepG2 cells. Henceforth, for simplicity, we will denote all the responses that involve substantial Trx-S^-^ depletion over a wide interval of intermediate *ν*_sup_ *(i.e.,* responses A, ADU, ADS, PDU and PDS from ST3) as “Response D”.

Fourth, the remaining 10 human cell lines are predicted to show broadly similar responses that are transitional between the generic responses P, PD, and PDS in ST3. Namely, they share the following features. (i) Little Trx-S^-^ depletion at medium-high values of *ν*_sup_. (ii) High Prx-SS accumulation at only a narrow range of *ν*_sup_, followed by progressively decreasing concentrations with increasing *ν*_sup_. (iii) An ultrasensitive increase in the cytoplasmic concentration of H_2_O_2_ and Prx-SO_2_^-^, and decrease of that of Prx-S^-^ by several orders of magnitude over this narrow *ν*_sup_ range. Henceforth we will denote this overall behavior as “Response H”.

If only 3% of the Trx1 is available to reduce Prx-SS, all the nucleated human cells are predicted to exhibit Response PD from ST3, where region DTPU appears between regions TTPU and HTAU (SF19). Accordingly, Prx-SS becomes the dominant Prx form over a range of *ν*_sup_ and the increase of Prx-SO_2_^-^ with *ν*_sup_ becomes more gradual (SF20). A comparison of Figure 3 and Figure 4 to SF19 and SF20 (respectively) highlights the following three points. First, a strong Trx oxidation can be prevented by decreasing the concentration of Trx available to reduce Prx-SS. Second, the decreased Trx availability shifts the *ν*_sup_ at which Prx-SS becomes dominant 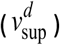 to lower values. Third, although the strongly decreased Trx availability makes the phenotype of all nucleated cells qualitatively similar, it increases the variability in cells tolerance to Prx oxidation.

### Tracing phenotype to molecular properties

What specific differences and similarities in protein composition underlie the differences and similarities in the predicted PTTRS phenotypes of human cells?

Response D and the high resistance of PrxII in human erythrocytes to hyperoxidation were attributed to the low TrxR activity in these cells [1]. The analysis below supports this view and provides a more complete understanding of the multiple factors underlying this outcome. It follows from rearranging the boundary conditions in ST2 that for the PTTRS to enter the regions where both Prx and Trx are mostly oxidized to disulfides (DDAS or DDAU) at some range of *ν*_sup_ the following condition must hold:

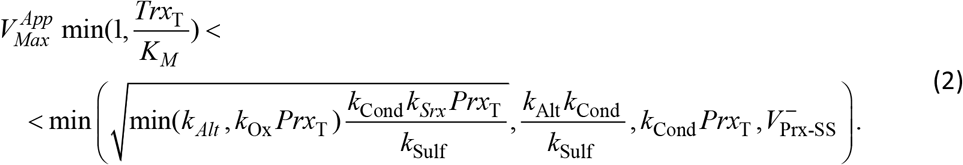

Here

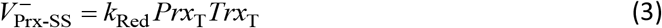

stands for the maximum rate of Prx-SS reduction when not limited by the TrxR activity. For the kinetic parameters and concentration ranges of Prx and Trx in most human cells in this study, 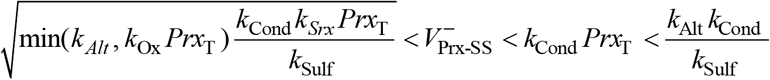 holds, and therefore condition (2) simplifies to:

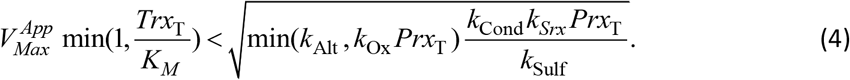

The left hand side of this inequality approximates the maximal possible rate of Trx-SS reduction in the system, and the right hand side approximates the maximal rate of Prx-SS production (and thus Trx oxidation) taking into account the balance between hyperoxidation and Prx-SO_2_^-^ reduction. Therefore, this inequality shows that a maximal Trx reduction rate lower than the maximal Trx oxidation rate warrants the accumulation of both Prx-SS and Trx-SS at some range of *ν*_Sup_.

In cells showing Response D the value of 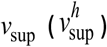 at which Prx-SO_2_^-^ becomes the dominant Prx form (*i.e*., the PTTRS crosses from regions DDAU or DDAS into region HTAU) is approximately:

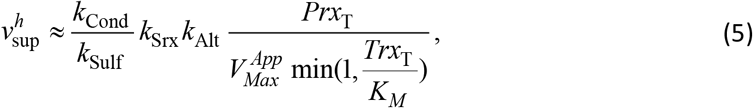

as can be derived from the expressions in ST2. Therefore, a high resistance of Prx to hyperoxidation in these cells depends on the following factors. (i) The intrinsic resistance of the dominant Prx to hyperoxidation (*k*_Cond_/*k*_Sulf_); (ii) a high Srx activity; (iii) the availability of alternative H_2_O_2_ sinks (*k*_Alt_); and (iv) a low Prx reduction turnover, as expressed by the last factor in expression (5). (The value of 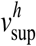 for erythrocytes inferred from Figure 3E is underestimated because Srx reduces PrxII more rapidly than PrxI [60] and because the PrxII-SO^-^ glutathionylation rate approaches the condensation rate,[42] thereby inhibiting hyperoxidation to some extent.)

In turn, the value of 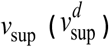 at which Prx-SS and Trx-SS become the dominant forms (*i.e.*, the PTTRS crosses from regions TTPU or TTAU into regions DDAU or DDAS) is approximately:

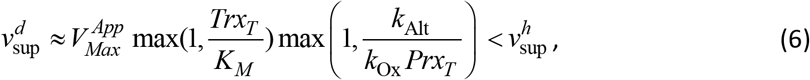

which for cells transitioning from TTPU into DDAS simplifies to:

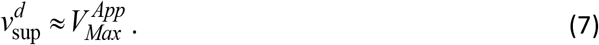

Response H emerges in cells where at some intermediate range of *ν*_sup_ the PTTRS crosses the overlap between regions TTPU, HTPU and HTAU. According to the design space analysis the range of *ν*_sup_ where the overlap occurs is determined by:

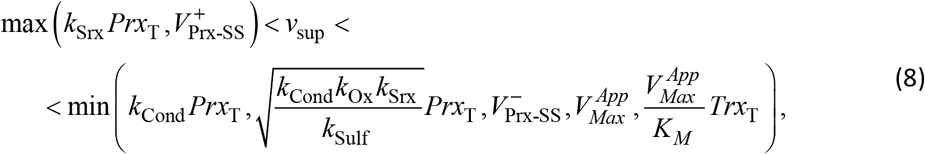

with

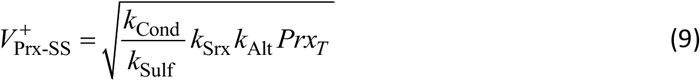

representing the maximal steady state rate of Prx oxidation to Prx-SS. This is a particular case of the expression in the right hand side of Expression (4) reflecting the fact that the overlap only occurs where *K*_Ox_*PrX_T_* > *k*_Alt_. Taking the kinetic parameters and protein concentration ranges in the human cell lines into account, the overlap range simplifies to:

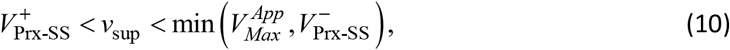

which yields the following condition for occurrence of the overlap:

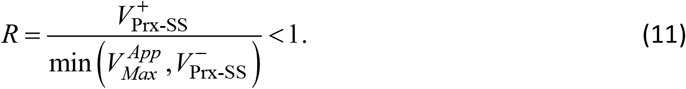

Note that the denominator approximates the maximal rate of Prx-SS reduction, and therefore this expression means that the overlap occurs when this maximal rate exceeds the maximal rate of Prx-SS production. With A549 cells as the only significant exception, in the examined nucleated human cells it is the TrxR activity that limits the rate of Prx-SS reduction, in absence of substantial Trx1 sequestration.

The overlap between regions indicates that multistability might occur, but the approximations in the design space analysis tend to underestimate the minimum and overestimate the maximum values of *ν*_sup_ for overlap (Figure 2C). As result, only Jurkat T cells are numerically predicted to show hysteresis over a very narrow *ν*_sup_ range (Figure 4F). But remarkably all Response H cells with known morphometry have *R* values in the range 0.61 to 0.76 (except *R*=0.34 for LnCap cells as an outlier), indicating that their PTTRS operates at the margins of the overlap region. For progressively lower values of this ratio the switch from low to high hyperoxidation becomes increasingly ultrasensitive and then hysteretic. Importantly, the maximum fraction of Prx in Prx-SS form and the maximum fractions of oxidized Trx attained at the switch get progressively lower.

For cells predicted to show Response H, Prx-SO_2_^-^ becomes the dominant Prx form at

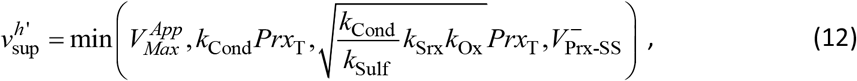

which in most cells corresponds to

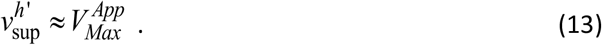

*R* is higher than 1 when 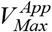 or *Trx_T_* are too low to cope with the maximum Prx-SS production rate. When the low 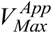 is the culprit, Response D ensues, as discussed above. In turn, when 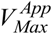 is high enough but *Trx_T_* is not, Prx can accumulate as Prx-SS without substantial Trx oxidation over a range of *ν*_sup_ (Region DTAU). This will happen in virtually all nucleated human cells in our set if only 3% of the *Trx_T_* estimated from the proteomic datasets is available to reduce Prx-SS as estimated above for HEK293 cells (SF19,20).

More precise conditions for occurrence of this behavior can be derived by rearranging the boundary conditions for region DTAU in ST2:

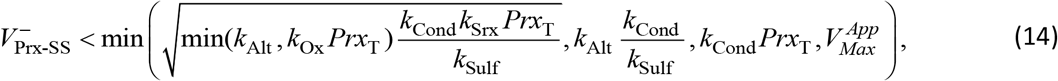

which for the kinetic parameters and protein concentration ranges in the human cell lines in this study simplifies to

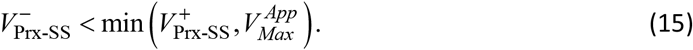

Under these conditions, Prx-SO_2_^-^ becomes the dominant Prx at [compare to (5)]

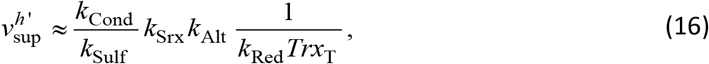

reflecting the fact that the Prx reduction turnover is then determined by Trx availability. In turn, Prx-SS becomes the dominant form at

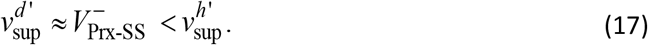

As *R* approaches 1, 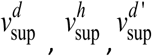 and 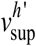 converge at

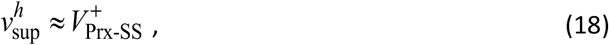

which is the minimum value of *ν*_sup_ at which Prx-SO_2_^-^ can become the dominant Prx form (Figure 2C).

### Enzyme activities in the PTTRS are correlated over cell lines

Is the similarity of the responses among human cell lines due to similar concentrations of the PTTRS proteins in all cells, or to correlations among the protein concentrations over cells? The following two observations support the latter possibility. First, Table 2 shows that there is substantial compositional heterogeneity among cell lines. For instance, *k*_Srx_ varies over a ≈20-fold range over cells showing Response H. Second, the values of *k*_Srx_ over these cells are very strongly correlated with the *k*_Srx_ threshold value,

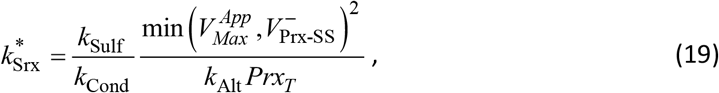

for occurrence of the overlap (Figure 5A).

**Figure 5.**
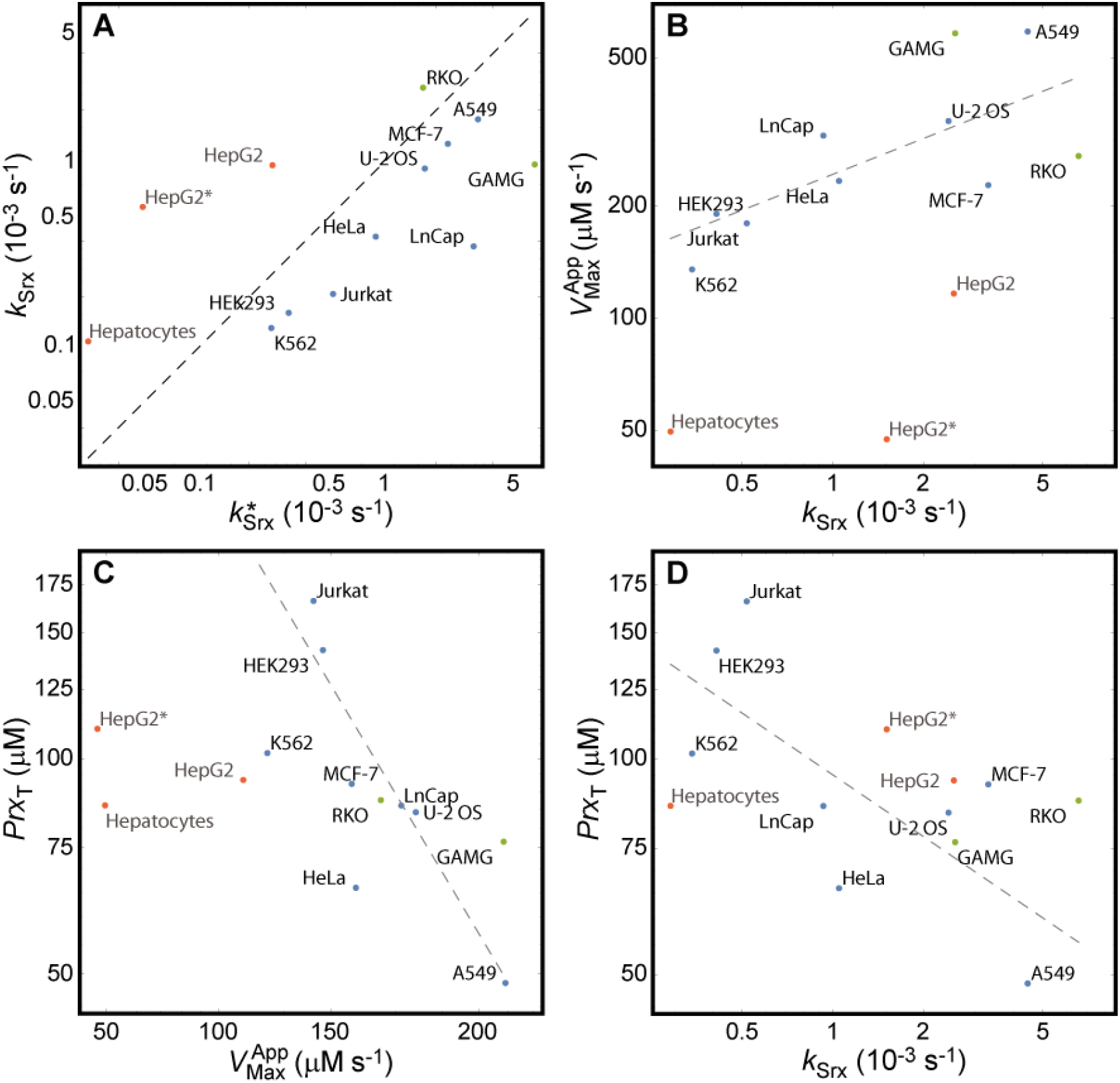
Correlations between parameters over cell lines. (A) k_Srx_ strongly correlates with the minimum value 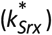 necessary to avoid the overlap region: Spearman rank correlation (σ_S_) 0.89, (p=0.0068) for all cell lines with Response H and known morphometry. Black dashed line indicates 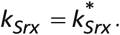 Red dots, cells showing Response D; green, cells with unknown morphometry showing response H; blue, cells with known morphometry showing response H. The following parameters are strongly correlated over cells with known morphometry showing Response H. (B) The activities of TrxR and Srx: σ_S_= 0.81, p= 0.015. Dashed line: best log-log fit over Response H cells with known morphometry, yielding a scaling exponent 0.38±0.13 (R^2^= 0.66). (C) The activity of TrxR and the total concentration of PrxI+PrxII: σ_S_= −0.86, p= 0.0065. Dashed line: best log-log fit over Response H cells with known morphometry, yielding a scaling exponent -0.87±0.31 (R^2^= 0.57). (D) The activity of Srx and the total concentration of PrxI+PrxII: σ_S_= −0.74, p= 0.037. Dashed line: best log-log fit over Response H cells with known morphometry, yielding a scaling exponent -0.29±0.11 (R^2^= 0.51). Note the logarithmic scales.

A statistical analysis reveals three prominent features of the orchestration among the parameters of the PTTRS over cells predicted to show Response H. First, a positive correlation between 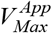 and 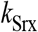 (Figure 5B). Remarkably, 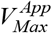 scales approximately as 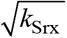 over these cell lines (Figure 5B), suggesting that the relative concentrations of TrxR and Srx are balanced in such way as to make the ratio between the maximal Prx-SS production rate and the maximal Trx-SS reduction rate approximately invariant over cell lines.

Second, a negative correlation between *k*_Srx_ and *PrX_T_* (Figure 5D). In Region TTPU, where most cells operate in absence of stress, the concentration of Prx-SO_2_^-^ is inversely proportional to the product *k*_Srx_*Prx_T_* (ST2). Furthermore, the value of *ν*_sup_ at which Prx-SO_2_^-^ becomes the dominant Prx form in Response H cells is approximately proportional to 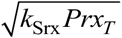. Therefore, the negative correlation between *k*_Srx_ and *Prx_T_* attenuates the heterogeneity of Prx-SO_2_^-^ concentration and hyperoxidation resistance among cell lines.

Third, a strong negative correlation between 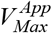 and *PtX_T_* (Figure 5C). This correlation may be just a consequence of the previous ones, as it lacks an obvious direct functional relevance.

Interestingly, 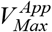 correlates with 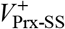 over Response H cells (σs= 0.65, p= 0.043), but *Trx_T_* and 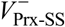 do not (σs= 0.28, p= 0.42; σs= 0.018, p= 0.96, respectively). This is in keeping with our estimates showing that Prx-SS reduction is rate limited by TrxR activity and not by Trx1 availability in the examined human cells, with A549 cells as the only significant exception. It also further supports the notions that Prx-SS reduction is the process with the greatest capacity to drive Trx1 oxidation (SI3.2.7) and that Trx1 is not strongly sequestered in most cells.

Finally, Figure 5 also highlights that Response D cells in our dataset have Srx and Prx concentrations in the range of the other cells but have disproportionally low TrxR.

### The PTTRS in yeast has distinctive properties

Does the dynamics of the PTTRS in other eukaryotic organisms differ from that in human cells? In order to address this question we estimated the parameters for *S. cerevisiae* (SI3.4). The design space analysis (Figure 3L) and numerical simulations of Model 1 (Figure 4L) based on these parameters predict that the responses of the PTTRS in the yeast depart from those described above for human cells in at least the following three aspects. First, Prx-SO^-^ concentrations are ≈7-fold higher than those of Prx-SS over all the range of *ν*_sup_, whereas in human cells Prx-SO^-^ concentrations are roughly similar to those of Prx-SS. Second, at intermediate values of *ν*_sup_ Prx accumulates mainly as Prx-SO^-^, and this remains the dominant Prx species over a wide range of *ν*_sup_ values. Third, the concentration of Prx-SO_2_^-^ increases more gradually with *ν*_sup_ than in human cells.

Table 2 shows many differences in protein composition and kinetic parameters between yeast and human cells. Which of these determine the differences between the behavior of the PTTRS in the yeast and in human cells? The high Prx-SO^-^/Prx-SS ratio in the yeast is explained as follows. The results in ST1 show that in all the phenotypic regions that can be attained in cells with high TrxR activity *(i.e.,* the *T*U regions) the ratio between the concentrations of Prx-SO^-^ and Prx-SS is approximated by *k*_Red_*TrX_T_* /*k*_Cond_. The about one order of magnitude lower cytoplasmic Trx concentration in the yeast is compensated by a ≈6-fold higher *k*_Red_ (SI3.4.3), yielding a similar value of *k*_Red_*Trx_T_*. However, the value of *k*_Cond_ is >10-fold lower for yeast’s Tsa1 than for human PrxI, explaining the higher ratio.

In turn, the following conditions for Prx-SO^-^ to become the dominant Prx species over a range of *ν*_sup_ can be derived from the results in ST2 for region STAU:

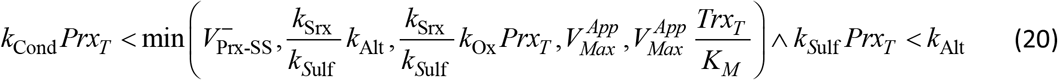

Both yeast and human cells fulfill the condition *k*_*S*ulf_*PrX_T_* <*k*_Alt_. However, only yeast cells fulfill the condition 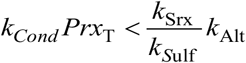. This is due to a combination of factors: (a) both *k*_Cond_ and *Prx*_T_ are lower in the yeast; (b) although both *k*_srx_ and *k*_Alt_ are also lower in the yeast, *k*_Sulf_ is much lower than in the human cells, making 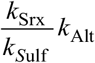 higher in the yeast. Therefore this behavior can be ascribed to the much higher stability of Tsa1’s sulfenate relative to human PrxI/II-SO^-^, and to the low Tsa1+Tsa2 concentration in the yeast.

The more gradual increase in Trx hyperoxidation in the yeast is explained by the following considerations. In the intermediate region STAU the concentration of Prx-SO^-^ is practically constant with *ν*_sup_, as most of the Prx was accumulated in this form. In turn, the concentration of H_2_O_2_ increases linearly with *ν*_sup_. As a consequence, the rate of hyperoxidation increases linearly with *ν*_sup_, rather than quadratically as in region TTPU.

## Discussion

### The PTTRS can respond to H_2_O_2_ supply in multiple ways

The results provide an approximated but simple description of the responses of the PTTRS in the cytoplasm of eukaryotic cells to physiological H_2_O_2_ supply rates (*ν*_sup_), and of how these responses depend on protein concentration and kinetic parameters. The approximations were thoroughly justified in the previous sections and in SI3, and are valid for *ν*_sup_ values that are insufficient to strongly deplete NADPH. For simplicity the model neglected the strong cytoplasmic concentration gradients of H_2_O_2_, Prx-SO^-^ and Prx-SS expected under low *ν*_sup_ [13].

The concentrations discussed below are thus spatially-averaged cytoplasmic concentrations. The analysis highlights the versatility of the PTTRS. For physiologically plausible parameters the PTTRS can exhibit eight qualitatively distinct types of steady states (“phenotypes”), corresponding to distinct hierarchies among fluxes and concentrations of the various Prx and Trx forms (Table 1, Figure 2, ST1). Two of these phenotypes allow the maintenance of Prx and Trx mostly in thiolate form and of strong concentration gradients. The remaining six are stress phenotypes where Prx is predominantly oxidized and gradients collapse.

The PTTRS can transition across 10 qualitatively distinct sequences of phenotypes as *ν*_sup_ increases from very low to very high values, defining as many distinct responses. In the process, the PTTRS has the potential to generate, proportional, saturable, ultrasensitive, non-monotonic, and even hysteretic behaviors. Cells may in principle take advantage of this diversity of behaviors to achieve distinct modes of regulation of downstream processes by H_2_O_2_ supply rate signals.

We mapped systematically the relationships among kinetic parameters and protein concentrations that make each of these phenotypes and responses emerge (ST2,3), and we derived simple approximations for the main PTTRS response thresholds. These phenotypes and thresholds usually depend on an interplay among multiple factors, as is well illustrated by the sensitivity to hyperoxidation *in vivo.* As shown by expressions (5) and (16) this property depends not only on the Prx’s intrinsic susceptibility of the Prx and on the Srx activity but also on the activity of non-Prx H_2_O_2_ sinks, on the Prx concentration and on the Prx-SS reduction capacity of the cytoplasm. Proper interpretation of the functional significance of gene expression, protein abundance or metabolic changes must take this interplay into account, which is difficult to accomplish without the help of modeling approaches such as illustrated here.

### The quantitative model is consistent with experimental observations and yields novel insights

In order to examine what of the potential behaviors discussed above can be realized in real cells, we estimated the necessary protein concentrations and kinetic parameters for ten human cancer cell lines, one non-cancer human cell line (HEK293), two differentiated human cell types (hepatocytes and erythrocytes), and *S. cerevisiae.* These estimates are based on quantitative proteomics data and published kinetic data. They are also thoroughly documented in SI3 and summarized in Table 2 and ST6, which we hope will provide a useful reference for quantitative redox biology.

With these data we first tested if simulations based on the parameterized models yielded results in agreement with available quantitative experimental data for human erythrocytes, Jurkat T cells and HEK293 cells. Computational predictions for the first two cell types were in very good agreement with the experimental observations (SF14,15).[1] Of note, they fully capture the observations that whereas treatment of 5×10^6^ erythrocytes/mL with H_2_O_2_ boluses up to 200 μM caused PrxII to accumulate in disulfide form and little Prx hyperoxidation, treatment of 10^6^ Jurkat T cells/mL with 100 μM or 200 μM H_2_O_2_ boluses caused extensive hyperoxidation [1].

In turn, the simulations for HEK293 cells agree qualitatively with the observations in ref. [18], but predict much lower Prx-SS fractions for the same extracellular H_2_O_2_ concentrations. This discrepancy is eliminated if only ≈3% (1.5 μM) of the Trx1 concentration estimated from the proteomic dataset is available to reduce Prx-SS (SF10A). Because the Trx1 concentrations Table 2 are in the range determined by other methods (see SI3.2.6) [61, 62], the most likely explanation for the discrepancy is the sequestration of Trx1 in the nucleus [63, 64] and/or in complexes with other proteins [65].

The following observations suggest that a larger fraction of Trx1 is available for Prx-SS reduction in other cell lines. First, the small fraction of PrxII oxidized to PrxII-SS in Jurkat T cells exposed to various H_2_O_2_ boluses in ref. [1] is not consistent with extensive Trx1 sequestration (SF14). Second, the TrxR activity, and not the Trx1 concentration, correlates with the maximum steady state rate of Prx-SS production over Response H cells, which suggests that in most cells Prx-SS reduction is limited by TrxR activity and not by Trx1 availability.

### Threshold behavior of Prx hyperoxidation and cytoplasmic H_2_O_2_ is intrinsic to the PTTRS dynamics

Surprisingly, this simple model for HEK293 cells (with *Trx*_T_ = 1.5 μM) shows very good agreement with the experimental observations [19] of a threshold H_2_O_2_ bolus beyond which Prx hyperoxidation starts increasing, and intracellular H_2_O_2_ increases more steeply (SF10B,C). Because the model considers neither a buffering effect by generic protein thiols nor saturable alternative H_2_O_2_ sinks, the observed agreement strongly argues against inferring the occurrence of such phenomena from the observed threshold behavior. Moreover, the balance of experimental evidence does not support the notion that generic (non-redoxin) protein thiols can provide a significant buffering capacity (see SI3.2.2.3). Instead, our results (SF11) indicate that the threshold behavior described in ref. [19] is intrinsic to the dynamics of the PTTRS. Concretely, the steeper increase in cytoplasmic H_2_O_2_ concentration and hyperoxidation starts once the bolus becomes sufficient to fully oxidize Prx to Prx-SS, thereby sharply decreasing the cytoplasmic H_2_O_2_ clearance rate. The cytoplasmic H_2_O_2_ concentration then becomes sufficient to cause accumulation of Prx-SO_2_^-^. Further supporting this interpretation, simulations using Model 1 to analyze the influence of the Trx concentration, TrxR activity, activity of alternative H_2_O_2_ sinks and Prx concentration on the threshold (SF12) agree with the observations in ref. [19]. There is a close correspondence between these thresholds and the *ν*_sup_ threshold marking the border of the TTPU region in the design space, whose physiological significance we will discuss below. Namely, they all reflect the saturation of cells’ Prx-SS reduction capacity.

### Human cell lines are predicted to show similar responses to H_2_O_2_ supply

Once applied to the quantitative estimates, the phenotypic map we derived allowed to make quantitative predictions about the responses of the PTTRS to H_2_O_2_ supply and TrxR modulation and to classify distinct cell types according to the predicted behavior of their PTTRS. This analysis revealed several intriguing commonalities and differences, which we discuss below.

When all the Trx1 is available to reduce Prx-SS in the cytoplasm, the 13 human cell types in our sample are predicted show just two major types of responses to *ν*_sup_ despite the diversity of cell types (ST4), and the variability in protein concentrations (ST6). Namely, in “Response D” intermediate values of *ν*_sup_ lead to oxidation of both Prx and Trx to disulfide forms, as well as to Trx-S^-^ depletion. In this response substantial hyperoxidation occurs only at very high *ν*_sup_. In turn, in “Response H” there is modest Trx-S^-^ depletion at any *ν*_sup_, and the steady state fraction of Prx-SS only reaches high values at a very narrow range of *ν*_sup_. In contrast, responses including an extensive accumulation of Prx-SS or Prx-SO^-^ without Trx-S^-^ depletion over a wide range of *ν*_sup_, or an abrupt bi-stable switch between a low and a high hyperoxidation state are possible (Figure 2, ST3) but not predicted to occur in any of the cells considered. (Jurkat T cells may show bistability over a tiny *ν*_sup_ range but this is strongly dependent on parameter uncertainties.)

Remarkably, all but one of the cell lines are predicted to exhibit Response H, and the two differentiated human cell types are predicted to exhibit Response D. The exception is the hepatoma HepG2 cell line. The model parameterizations based on both independent proteomic datasets for HepG2 [53, 54] consistently lead to the prediction that these cells exhibit the same type of response as hepatocytes: Response D. This observation raises the question of whether cancer cell lines tend to respond to *ν*_sup_ similarly to the differentiated cell types they derive from, but unfortunately we are unaware of other data sets that would permit such a comparison for other cell types.

Our analysis revealed the underpinnings of the predicted similarities and distinctions among the PTTRS responses of human cells. Response D occurs in cells where the maximal Trx reduction rate is substantially lower than the maximal Trx-SS production rate [Expression (4)]. Response H occurs where the TrxR activity and/or the maximal Prx-SS reduction slightly exceed the maximal Prx-SS production rate [0.3 ≤ *R* ≤ 1 in expression (11)].

In turn, when just ≈3% of the Trx1 is considered available to reduce Prx-SS, all *nucleated* human cells in our sample are predicted to show Response PD (ST3), which differs from Response H in that Prx-SS becomes the predominant Prx form over a wide range of *ν*_sup_ (SF19,20). The threshold value of *ν*_sup_ at which the Prx-S^-^ pool collapses also decreases. While for Response D and for most cells showing Response H this threshold occurs at 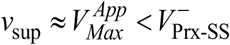 [expressions (7) and (13)], for Response PD it occurs at 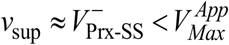 [expression (17)].

While the conditions above for occurrence of each type of response are intuitive and nearly trivial when stated in terms of maximum steady state fluxes, these fluxes have a more complex dependence on the kinetic parameters and protein concentrations. However, the design space approach permitted deriving relatively simple approximations that can be straightforwardly used to predict responses and threshold values.

### Landmarks of the quantitative design space have a physiological correspondence

Irrespective of the predicted Response type, under physiological, non-stress conditions all human cells analyzed here operate far within regions TTAU (erythrocytes) or TTPU (all others). As *ν*_sup_ increases, the fraction of Prx in oxidized forms gradually increases. Then, as *ν*_sup_ approaches 50–500 μM s^-1^, corresponding to 5–50 μM extracellular H_2_O_2_, nucleated human cells transition into regions DTAU, DDAS or HTAU, where Prx-SS or Prx-SO_2_^-^ predominate (Figure 3,Figure 4,SF17,18). This transition is accompanied by an ultrasensitive increase of the intracellular H_2_O_2_ concentration from the 1–10 nM to the μM range over a relatively narrow range of *ν*_sup_ (Figure 4, SF18). The sharp increase is due to cellular H_2_O_2_ sinks other than PrxI and PrxII being less active towards H_2_O_2_ (PrxVI), much less abundant (GPx1) and/or not directly accessible from the cytoplasm (Cat). The inability of these sinks to maintain a strong transmembrane H_2_O_2_ gradient is experimentally demonstrated by the observation that partial Cat inhibition has a substantial effect on the pseudo-first order rate constant (*k_cells_*) for the consumption of extracellular H_2_O_2_ boluses that are sufficient to extensively oxidize PrxI and PrxII [66, 67]: otherwise H_2_O_2_ consumption would be strongly limited by the membrane permeation step and virtually insensitive to Cat inhibition (see SI3.2.1). It is also supported by recent experiments in living zebra fish.[59] In turn, a strong transmembrane gradient when Prx-S^-^ is predominant is predicted by reaction-diffusion models [13, 68] and was experimentally estimated as ≈650 for HeLa cells [68].

Could the Prx-S^-^ collapse and ensuing sharp increase in cytoplasmic H_2_O_2_ associated to the transition from TTPU to the stress regions mark the threshold for H_2_O_2_ toxicity? Antunes & Cadenas [69] found that steady state H_2_O_2_ induces apoptosis of Jurkat T cells only above a threshold extracellular concentration. Moreover, this threshold is very sharp, such that doubling the H_2_O_2_ concentration from the threshold value causes the fraction of apoptosing cells to increase from control to saturation values. The threshold is also dependent on exposure duration, suggesting that H_2_O_2_ toxicity manifests only beyond a given dosage (concentration×time) threshold.[69] Huang & Sikes [68] recently reported similar findings for HeLa cells with respect to intracellularly produced H_2_O_2_ and further elaborated this concept. Interestingly, the extracellular H_2_O_2_ concentration where the TTPU→HTAU transition in Jurkat T cells is expected (Figure 3F, Figure 4F) corresponds approximately to the steady state concentration observed to cause apoptosis of 50% of these cells [69]. Furthermore, steady state extracellular H_2_O_2_ concentrations that are just enough to fully oxidize PrxII to PrxII-SS, but not lower concentrations, cause apoptosis of HEK239 cells.[18] According to our simulations of the PTTRS in these cells (SF13), under these circumstances PrxI should also become fully oxidized and intracellular H_2_O_2_ should attain the μM range. Therefore, these observations support the notion that the onset of H_2_O_2_ toxicity is associated to the collapse of the Prx-S^-^ pool. Accordingly, it should be modulated by the TrxR activity in cells showing Responses H or D, or Trx1 availability in cells where extensive Trx1 sequestration dictates Response PD. These predictions are experimentally testable.

The considerations above question the assertion that perturbations of the intracellular H_2_O_2_ concentration in the order of 10 nM can induce apoptosis. This assertion was based on a re-evaluation of the transmembrane H_2_O_2_ gradient to account for the activity of the cytoplasmic Prx.[68] This estimated ≈650-fold transmembrane gradient applies where the H_2_O_2_ supply rate to the cytoplasm is insufficient to strongly oxidize Prx *(i.e.,* well within region TTPU). However, apoptosis seems to be induced only upon the collapse of the Prx-S^-^ pool. Under these conditions the transmembrane gradient should be no higher than the ≈7-fold determined in ref. [70], and intracellular H_2_O_2_ concentrations should then approach the μM range [69].

Could the changes in Prx and/or Trx redox state that occur as the PTTRS transitions from region TTPU to the stress regions directly regulate apoptosis? Apoptosis induction by H_2_O_2_ is mediated by activation of apoptosis signal-regulating kinase 1 (ASK1). In the most consensual redox regulation model, this protein is inhibited by non-covalent binding of Trx1-S^-^ to its Trx1-binding domain, and released from inhibition by Trx1-S^-^ oxidation to Trx-SS.[71] However, the dissociation constant for the complexes of Trx1-S^-^ and Trx1-SS with ASK1’s Trx1-binding domain — 0.3±0.1 μM [72] and 4±2 μM [73], respectively — are *both* much lower than typical cytoplasmic Trx1 concentrations (Table 2). Therefore, not even the full oxidation of Trx can dissociate the inhibitory Trx1:ASK1 complex unless the dissociation from Trx1-SS is coadjuvated by other factors. These might include destabilizing interactions with other proteins (e.g. tumor necrosis factor receptor-associated factors 2 and 6), extensive Trx1 sequestration and/or oxidation of ASK1’s Cys residues.[74] The latter might be driven by the recently characterized redox relay mediated by Prx1-SS.[9] It is thus tempting to speculate that the PrxI-SS accumulation as cells cross from phenotypic region TTPU to DTAU, DDAS or HTAU might thereby trigger apoptosis. However, ASK1 regulation is complex and still poorly understood [75], and none of the proposed models explains, for instance, the iron dependence of H_2_O_2_-induced apoptosis [69].

Although if sustained over a long time a strong oxidation of the Prx-S^-^ pool may trigger cell death, temporary strong Prx oxidation does occur in the normal physiology of higher organisms. The events associated to zebra fish wounding illustrate this point: 2-5 μM extracellular H_2_O_2_ concentrations are attained within ≈30 nm of wound margins during tens of minutes, which strongly oxidize the cells’ Prx-S^-^ pool.[59] Studying the steady state response to H_2_O_2_ supply rates beyond region TTPU yields insight on cells’ responses to such temporary H_2_O_2_ surges. In all human cells examined in this work, sudden exposure to *ν*_sup_ beyond the threshold initially causes a short-lived surge in Prx-SO^-^ and then extensively oxidizes Prx-S^-^ to Prx-SS, dramatically increasing the cytoplasmic H_2_O_2_ concentration within seconds (SF21). In Response H and Response D cells, but not in Response PD cells, Trx-S^-^ is also rapidly and extensively oxidized. In turn, Prx-SO_2_^-^ accumulates much slower. In Response H cells, Prx-SO_2_^-^ may eventually accumulate as the dominant Prx form thereby relieving the load on the Trx-S^-^ pool. However, in other cells this would only happen at *ν*_sup_ beyond the thresholds defined by expressions (5), (16) which cells are unlikely to survive. Recovery of the Prx-S^-^ pool from Prx-SS once the stimulus stops can occur in seconds, but recovery from Prx-SO_2_^-^ can take hours.

### Dynamics of the PTTRS defines two distinct H_2_O_2_ signaling regimes

The considerations above have important implications for redox signaling in the cytoplasm of eukaryotic cells. The much higher activity of PrxI and PrxII as H_2_O_2_ reductants relative to the alternative H_2_O_2_ sinks determines the existence of two very distinct H_2_O_2_ signaling regimes with physiological relevance. For *ν*_sup_ below the cytoplasmic capacity to reduce Prx-SS, cytoplasmic H_2_O_2_ concentrations are determined by the Prx-S^-^ pool. Under these conditions there are strong H_2_O_2_ concentration gradients both across the cell membrane and over the cytoplasm.[13, 68] And even though cytoplasmic H_2_O_2_ concentrations are much higher near H_2_O_2_ supply sites than elsewhere, they are far too low to directly oxidize thiolates other than those in the active sites of peroxiredoxins and peroxidases in a signaling time frame even there.[13] Therefore, in this regime Prx must act as primary H_2_O_2_ sensors, which then actuate regulatory targets through localized[13] redox relays [9,10,76–78] or non-covalent binding/release of the oxidized Prx forms [11]. H_2_O_2_ chemotaxis by leukocytes far away from wound borders may be an example of H_2_O_2_ signaling in this regime.[59]

In turn, at *ν*_sup_ above the cytoplasmic capacity to reduce Prx-SS, cytoplasmic H_2_O_2_ concentrations are determined by the activities of peroxiredoxin VI, GPx1 and Cat. These activities are, collectively, too low to impose a significant H_2_O_2_ gradient over the cytoplasm or even a very large transmembrane gradient. Thus, H_2_O_2_ concentrations are expected to be in the μM range throughout the cytoplasm. However, as per the previous section, cells are unlikely to survive such H_2_O_2_ concentrations for more than a few tens of minutes, which constrains the redox targets that can be directly actuated by H_2_O_2_ even in this regime: For 50% of a target’s molecules to be oxidized within 1h by 1 μM H_2_O_2_ the oxidation rate constant must exceed ln(2)/(3600 s × 10^−6^ M)= 193 M^-1^s^-1^. Indeed, several regulatory targets that are involved in adaptation to oxidative stress and thus should be actuated in this regime and not in the low-*ν*_sup_ one, have H_2_O_2_ reactivities in this range. These include glyceraldehyde 3-phosphate dehydrogenase (GAPDH, *k*= 500 M^-1^s^-1^ [79, 80]), Kelch-like ECH-associated protein 1 (KEAP1, *k*= 140 M^-1^s^-1^ [81]), and a still unidentified target that regulates nuclear factor erythroid 2-related factor 2 (NRF2) protein synthesis (*k*≥ 1300 M^-1^s^-1^ [81]). Of these, the oxidative inhibition of GAPDH by H_2_O_2_ contributes for adaptation by redirecting the metabolic flux from glycolysis to the oxidative part of the pentose phosphate pathway flux, thereby increasing NADPH regeneration.[82] The reactions with KEAP1 and the NRF2 regulator induce multiple antioxidant defenses whose promoters carry the antioxidant response element. On the other hand, various protein tyrosine phosphatases that are rapidly inactivated upon cells’ stimulation with mitogenic factors have substantially lower rate constants for direct oxidation by H_2_O_2_.[83, 84] Therefore, even in this regime their inactivation requires other mechanisms (e.g. peroximonocarbonate-mediated [85, 86]).

The analysis of a simpler reaction-diffusion model of H_2_O_2_ signaling also supports the notion of two distinct regimes,[13] with the transition here occurring through the hysteretic switch to the high-hyperoxidation regime (as in Figure 2C, bottom). This hysteretic switch is unlikely to occur in the cells examined in the present work, except under glutathione depletion (discussed below). However, this does not change the remaining conclusions in ref. [13]. In particular, the estimated times required to actuate redox targets by direct oxidation by H_2_O_2_ remain pertinent. It has been argued [86] that by neglecting target reduction ref. [13] overestimated the signaling response times. However, the analysis in ref. [86] refers to the time required for the target’s redox state to approach a new steady state, not to the time required to oxidize a given fraction of the target’s molecules as ref. [13] does. Target reduction cannot shorten the latter time. And it only accelerates the response time by decreasing the fraction of the target that is oxidized at the new steady state — that is, fewer target molecules need to be oxidized for approaching the new steady state. The experimental observations that most targets are substantially oxidized upon the stimuli indicate that reduction could accelerate the responses perhaps by up to 3-fold, but not by orders of magnitude.

### Is Response H an optimal trade-off between signaling and protection?

Why is Response H so prevalent among human cell lines? Could this be explained by any functional advantages? A main distinguishing feature of Response H over Responses D and PD is that at high *ν*_sup_ Prx-SO_2_^-^ gradually accumulates, eventually replacing Prx-SS as the dominant Prx form (SF21A). This behavior may have the following advantages.

First, it may better protect nascent proteins in proliferating cells against aggregation. The strong increase in cytoplasmic H_2_O_2_ to μM concentrations accompanying the collapse of the PTTRS’ redox capacity should dramatically accelerate oxidative damage by Fenton-reaction-derived hydroxyl radicals, lipid-peroxidation-derived electrophiles, etc. Recently synthesized proteins that have not yet folded [87] and misfolded proteins [88] are particularly vulnerable to this damage and to aggregation. Interestingly, hyperoxidation converts Prx to efficient holdases [89–91], which protect damaged and misfolded proteins against aggregation [92]. The following recent findings [93] support an even more central role of Prx-SO_2_^-^ in protection against H_2_O_2_-induced protein aggregation/inclusions. In *S. cerevisiae,* hyperoxidized Tsa1 recruits Hsp70 chaperones to damaged or unfolded proteins, thereby promoting their folding or destruction. Additionally, it recruits Hsp70/Hsp104 to H_2_O_2_-induced protein inclusions, promoting their disaggregation. Something similar may happen in human cells. Accordingly, PrxII hyperoxidation protected HeLa cells from H_2_O_2_-induced apoptosis, whereas hyperoxidation-resistant, peroxidase-competent PrxII mutants were less protective [90]. Supporting a connection with cell proliferation, PrxII hyperoxidation correlated with reversible H_2_O_2_-induced cell cycle arrest of C10 mouse lung epithelial cells [94].

Second, the higher hyperoxidation decreases Prx-SS production and the ensuing load on the Trx-S^-^ pool (compare SF21A to SF21G), which has at least three advantages. (A) Leaving this pool available for other protective processes such as the reduction of methionine sulfoxides.[95] (B) Improving resistance to apoptosis by delaying ASK1 activation. (C) Decreasing the excretion of pro-inflammatory glutathionylated PrxI-SS and PrxII-SS [96–98]. Excretion of these Prx forms is proportional to the intracellular Prx-SS concentration [97], and it may signal the imminent collapse of the cell’s ability to sustain oxidative stress.

Also important, in cells lacking sufficient TrxR activity or Trx1 to avoid Responses D and PD the Prx-S^-^ pool collapses at lower *ν*_sup_ than in cells with higher TrxR and Trx concentrations and otherwise identical protein concentrations and activities. Those cells may thus be less tolerant of H_2_O_2_ supply.

The discussion in the previous paragraphs raises the question of why shouldn’t cells carry more Trx1 and TrxR, as this might increase their resistance to H_2_O_2_. Perhaps the physiological H_2_O_2_ supply rates are rarely high enough for the advantages of increased resistance to compensate the costs of increased protein expression. But there may be two other reasons. First, higher Trx1 and TrxR would cause bi-stability, such that transient increases in *ν*_sup_ beyond 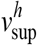 in Expression (18) could then trigger the positive feedback causing near-complete Prx hyperoxidation (Figure 2C, bottom). This positive feedback is intensified, making the switch more likely, if the GSH pool is depleted. This is both because the decrease in H_2_O_2_ clearance due to hyperoxidation then has a stronger impact on H_2_O_2_ concentrations and because Srx-catalyzed Prx-SO_2_^-^ reduction is itself GSH-dependent. Recovery from the high-hyperoxidation state can start only after *ν*_sup_ decreases below 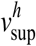 and is very slow, limited by the low Srx activity. Therefore, cells would be left with a diminished H_2_O_2_ clearance capacity for a long time, which would eventually trigger apoptosis. Intriguingly, the Trx1 and TrxR concentrations in almost all cell lines in our sample are just low enough to avoid this bi-stable behavior, which suggests that precisely controlling the extent of hyperoxidation is critical for optimal antioxidant protection.

The second reason to avoid very high Trx1 and TrxR concentrations is that this would prevent the attainment of high Prx-SS and/or Trx-SS concentrations, thereby disengaging redox relays mediated by these species.

Altogether, the considerations above suggest that Response H embodies an optimal trade-off between effective signal transduction and effective antioxidant protection. Together with the prediction that both differentiated human cell types in our sample show Response D they also raise the question of whether Response H is associated to cell proliferation and helps tumor cells survive.

The hypothesized functional advantages of Response H are experimentally testable through genetic or pharmacological manipulation of the PTTRS.

### Strong correlations among protein concentrations over cell lines ensure the maintenance of Response H

The predicted similarity among the responses of the PTTRS is not simply due to a similar protein composition of the cell lines. Instead, it is due to strong correlations among the concentrations of several proteins over cell lines in such a manner as to keep the maximum rate of Prx-SS production and the capacity for Prx-SS reduction approximately balanced. A positive correlation between 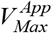 and *k*_Srx_ (Figure 5B), and negative correlations between 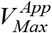 and *Prx_T_* (Figure 5C) as well as between *k_Srx_* and *Prx_T_* (Figure 5D) are strong and statistically significant. However, these three correlations are insufficient to explain the low variability of *R* over these cell lines, as can be verified by replacing the scalings described in Figure 5B-D into Expression (11). The additional dependencies among protein concentrations and activities that are required to explain the invariance are likely embedded in various other correlations that border the significance threshold. Thus, larger cell type samples and/or more precise determinations may reveal additional aspects of the orchestration among the concentrations of the PTTRS’ proteins.

This orchestration should be at least in part the result of developmental processes associated to cell differentiation and determining, for instance, that mean Srx concentrations are >100-fold higher in some human cell types than in others. However, it likely also reflects faster (hours-scale) regulatory mechanisms ensuring that each cell keeps a proper balance among the PTTRS’ proteins despite substantial fluctuations (“noise”) of each protein’s concentration. Microscopy techniques allowing to follow the abundances of, say, fluorescently labeled Srx and TrxR in single cells over time may clarify this point in the near future.

Given the strong involvement of the PTTRS in cancer and vascular processes [99–101], the characterization of these regulatory mechanisms may have important implications for human health. For instance, therapies that interfere with these mechanisms thereby unbalancing the PTTRS may be more effective against cancer than therapies modulating the activity of any single PTTRS protein.

The orchestration discussed above also implies that the functional consequences of up- or down-regulation of any single PTTRS protein must be evaluated in light of likely compensatory regulation of the other proteins. Quantifying multiple selected proteins will thus be important for understanding potential disturbances of the PTTRS in disease. However, even these multiplexed approaches may yield little insight in lack of a proper theoretical framework to help interpret the results.

### Yeast show a distinctive response to H_2_O_2_ supply

The strong similarities in the response of the PTTRS across human cell lines prompted to question if the PTTRS from phylogenetically distant eukaryotes shows similar responses. This turns out not to be the case. *S. cerevisiae* was predicted to exhibit a distinctive response to H_2_O_2_ supply: its main Prx (Tsa1p) accumulates predominantly as Prx-SO^-^ over a wide range of intermediate *ν*_sup_ (Figure 3L, Figure 4L). Furthermore, over all values of *ν*_sup_ Prx-SO^-^ concentrations are ≈7-fold higher than those of Prx-SS, whereas in human cells they are similar or lower than those of Prx-SS. That is mainly a consequence of the low *k*_Cond_ = 0.58 s^-1^ estimated for Tsa1 (SI3.4.3) from the data in ref. [102]. The low *k*_Cond_ and consequent high Prx-SO^-^ concentrations might make the yeast very sensitive to hyperoxidation. But remarkably the same data indicate that Tsa1’s sulfenate group is exceptionally stable, with a *k*_Sulf_ = 7. M^-1^s^-1^ (estimated in SI3.4.3), as compared to 1.2 × 10^4^ M^-1^s^-1^ for human PrxII and PrxIII [103] and estimated 1.3 ×10^3^M^-1^s^-1^ for PrxI (estimated in SI3.2.3). Consequently, the yeast is predicted (Figure 4L) to be more resistant to Prx hyperoxidation than most human cell lines, which is indeed experimentally observed [91].

How these “special” properties of Tsa1 relate to its recently characterized [93] roles in proteostasis and aging is an interesting matter for future research.

## Concluding remarks

The relationship between the PTTRS’ responses to H_2_O_2_ and its proteins’ concentrations and properties is complex, but it can be expressed in terms of simple analytical approximations. The concentrations-to-phenotype map presented here can guide the interpretation of protein concentration or gene expression datasets as well as the design of new redox biology experiments and therapies.

The analysis identifies the Prx-SS reduction capacity as a critical factor separating two distinct H_2_O_2_ signaling regimes in all nucleated human cells. At H_2_O_2_ supply rates below Prx-SS reduction capacity, H_2_O_2_ concentrations are very low throughout the cytoplasm and highly localized. Signaling in this regime should thus be mediated by localized Prx-mediated redox relays. In turn, at H_2_O_2_ supply rates above Prx-SS reduction capacity μM-range H_2_O_2_ concentrations are attained uniformly throughout the cytoplasm. Signaling in this regime can occur through direct oxidation of some targets by H_2_O_2_. But as cells are unlikely to tolerate such high cytoplasmic H_2_O_2_ for more than a few tens of minutes this signaling mechanism is only effective for targets with a H_2_O_2_ reactivity >10^2^ M^-1^s^-1^.

Nearly all human cell lines analyzed are predicted to show a similar response to H_2_O_2_ supply that may optimize a trade-off between effective redox-relay signaling and effective antioxidant protection. This response favors a gradual and moderate accumulation of hyperoxidized Prx after the H_2_O_2_ supply exceeds the above-mentioned threshold. It hinges on the Prx-SS reduction capacity exceeding the maximal steady state Prx-SS production rate by a small margin that is insufficient to enable a hysteretic switch to a full-hyperoxidation state. This balance between maximal Prx-SS production and reduction capacity is maintained through strong correlations between the concentrations of TrxR, Srx, and Prx over cell lines.

The Tsa1-based PTTRS in *S. cerevisiae* is predicted to show a distinctive response to H_2_O_2_ supply, largely by virtue of the exceptional stability of Tsa1-SO^-^.

The phenotypic map can be gradually refined as the kinetic properties of further relevant redox interactions are determined, allowing to better define the responses to high H_2_O_2_ supply rates, the functional complementarity between PrxI and PrxII, and the interactions between the PTTRS and the Grx/GSH/GSSG reductase system.

## Acknowledgements

This work was funded by fellowship SFRH/BD/51576/2011 and grants UID/NEU/04539 COMPETE (POCI-01-0145-FEDER-007440), PEst-OE/QUI/UI0612/2013, and FCOMP-01-0124-FEDER-020978 financed by FEDER through the “Programa Operacional Factores de Competitividade, COMPETE” and by national funds through “FCT, Fundação para a Ciência e a Tecnologia” (project PTDC/QUI-BIQ/119657/2010). We thank Dr. Michael Savageau and Dr. Jason Lomnitz (University of California – Davis) for illuminating discussions, and Ms. Lisa Susi for careful review of the manuscript.

## References

[1] F.M. Low, M.B. Hampton, A. V Peskin, C.C. Winterbourn, Peroxiredoxin 2 functions as a noncatalytic scavenger of low-level hydrogen peroxide in the erythrocyte, Blood. 109 (2007) 2611–2617. doi:10.1182/blood-2006-09-048728.

[2] R.M. Johnson, J. r Goyettejr G, Y. Ravindranath, Y.S. Ho, Hemoglobin autoxidation and regulation of endogenous H2O2 levels in erythrocytes, Free Radical Biology and Medicine. 39 (2005) 1407–1417. doi:10.1016/j.freeradbiomed.2005.07.002.

[3] P.A. Karplus, A primer on peroxiredoxin biochemistry, Free Radical Biology and Medicine. 80 (2015) 183–190. doi:10.1016/j.freeradbiomed.2014.10.009.

[4] M.H. Choi, I.K. Lee, G.W. Kim, B.U. Kim, Y.H. Han, D.Y. Yu, H.S. Park, K.Y. Kim, J.S. Lee, C. Choi, Y.S. Bae, B.I. Lee, S.G. Rhee, S.W. Kang, Regulation of PDGF signalling and vascular remodelling by peroxiredoxin II, Nature. 435 (2005) 347–353. doi:10.1038/nature03587.

[5] H.A. Woo, S.H. Yim, D.H. Shin, D. Kang, D.Y. Yu, S.G. Rhee, Inactivation of Peroxiredoxin I by Phosphorylation Allows Localized H2O2 Accumulation for Cell Signaling, Cell. 140 (2010) 517–528. doi:10.1016/j.cell.2010.01.009.

[6] D.Y. Jin, H.Z. Chae, S.G. Rhee, K.T. Jeang, Regulatory Role for a Novel Human Thioredoxin Peroxidase in NF-κB Activation, The Journal of Biological Chemistry. 272 (1997) 30952–30961. doi:10.1074/jbc.272.49.30952.

[7] J. Cao, J. Schulte, A. Knight, N.R. Leslie, A. Zagozdzon, R. Bronson, Y. Manevich, C. Beeson, C.A. Neumann, Prdx1 inhibits tumorigenesis via regulating PTEN/AKT activity, The EMBO Journal. 28 (2009) 1505–1517. doi:10.1038/emboj.2009.101.

[8] S.Y. Kim, T.J. Kim, K.Y. Lee, A novel function of peroxiredoxin 1 (Prx-1) in apoptosis signal-regulating kinase 1 (ASK1)-mediated signaling pathway, FEBS Letters. 582 (2008) 1913–1918. doi:10.1016/j.febslet.2008.05.015.

[9] R.M. Jarvis, S.M. Hughes, E.C. Ledgerwood, Peroxiredoxin 1 functions as a signal peroxidase to receive, transduce, and transmit peroxide signals in mammalian cells, Free Radical Biology and Medicine. 53 (2012) 1522–1530. doi:10.1016/j.freeradbiomed.2012.08.001.

[10] C.C. Winterbourn, Reconciling the chemistry and biology of reactive oxygen species, Nature Chemical Biology. 4 (2008) 278–286. doi:10.1038/nchembio.85.

[11] C.A. Neumann, J. Cao, Y. Manevich, Peroxiredoxin 1 and its role in cell signaling, Cell Cycle. 8 (2009) 4072–4078. doi:10.4161/cc.8.24.10242.

[12] Z.A. Wood, Peroxiredoxin Evolution and the Regulation of Hydrogen Peroxide Signaling, Science. 300 (2003) 650–653. doi:10.1126/science.1080405.

[13] R.D.M. Travasso, F. Sampaio dos Aidos, A. Bayani, P. Abranches, A. Salvador, Localized Redox Relays as a Privileged Mode of Cytoplasmic Hydrogen Peroxide Signaling, Redox Biology. 12 (2017) 233–245. doi:10.1016/j.redox.2017.01.003.

[14] H.A. Woo, H.Z. Chae, S.C. Hwang, K.S. Yang, S.W. Kang, K. Kim, S.G. Rhee, Reversing the Inactivation of Peroxiredoxins Caused by Cysteine Sulfinic Acid Formation, Science. 300 (2003) 653–656. doi:10.1126/science.1080273.

[15] T. Geiger, A. Wehner, C. Schaab, J. Cox, M. Mann, Comparative proteomic analysis of eleven common cell lines reveals ubiquitous but varying expression of most proteins, Molecular & Cellular Proteomics: MCP. 11 (2012) M111.014050. doi:10.1074/mcp.M111.014050.

[16] J.H. Seo, J.C. Lim, D.Y. Lee, K.S. Kim, G. Piszczek, H.W. Nam, Y.S. Kim, T. Ahn, C.H. Yun, K. Kim, P.B. Chock, H.Z. Chae, Novel Protective Mechanism against Irreversible Hyperoxidation of Peroxiredoxin N-alpha-terminal acetylation of human peroxiredoxin, Journal Of Biological Chemistry. 284 (2009) 13455–13465. doi:10.1074/jbc.M900641200.

[17] A. Mitsumoto, Y. Takanezawa, K. Okawa, A. Iwamatsu, Y. Nakagawa, Variants of peroxiredoxins expression in response to hydroperoxide stress, Free Radical Biology and Medicine. 30 (2001) 625–635. doi:10.1016/S0891-5849(00)00503-7.

[18] M.C. Sobotta, A.G. Barata, U. Schmidt, S. Mueller, G. Millonig, T.P. Dick, Exposing cells to H2O2: A quantitative comparison between continuous low-dose and one-time high-dose treatments, Free Radical Biology and Medicine. 60 (2013) 325–335. doi:10.1016/j.freeradbiomed.2013.02.017.

[19] L.E. Tomalin, A.M. Day, Z.E. Underwood, G.R. Smith, P. Dalle Pezze, C. Rallis, W. Patel, B.C. Dickinson, J. Bähler, T.F. Brewer, C.J.-L. Chang, D.P. Shanley, E.A. Veal, Increasing extracellular H2O2 produces a bi-phasic response in intracellular H2O2, with peroxiredoxin hyperoxidation only triggered once the cellular H2O2-buffering capacity is overwhelmed, Free Radical Biology and Medicine. 95 (2016) 333–348. doi:10.1016/j.freeradbiomed.2016.02.035.

[20] C.S. Pillay, J.H. Hofmeyr, L.N. Mashamaite, J.M. Rohwer, From Top-Down to Bottom-Up: Computational Modeling Approaches for Cellular Redoxin Networks, Antioxidants & Redox Signaling. 18 (2013) 2075–2086. doi:10.1089/ars.2012.4771.

[21] R. Benfeitas, G. Selvaggio, F. Antunes, P.M.B.M. Coelho, A. Salvador, Hydrogen peroxide metabolism and sensing in human erythrocytes: A validated kinetic model and reappraisal of the role of peroxiredoxin II, Free Radical Biology and Medicine. 74 (2014) 35–49. doi:10.1016/j.freeradbiomed.2014.06.007.

[22] J.B. Lim, T.F. Langford, B.K. Huang, W.M. Deen, H.D. Sikes, A reaction-diffusion model of cytosolic hydrogen peroxide, Free Radical Biology and Medicine. 90 (2016) 85–90. doi:10.1016/j.freeradbiomed.2015.11.005.

[23] C.S. Pillay, B.D. Eagling, S.R. Driscoll, J.M. Rohwer, Quantitative measures for redox signaling, Free Radical Biology and Medicine. 96 (2016) 290–303. doi:10.1016/j.freeradbiomed.2016.04.199.

[24] A. Salvador, F. Antunes, R.E. Pinto, Kinetic modelling of in vitro lipid peroxidation experiments–’low level’ validation of a model of in vivo lipid peroxidation, Free Radical Research. 23 (1995) 151–172. doi:0.3109/10715769509064029.

[25] F. Antunes, A. Salvador, H.S. Marinho, R. Alves, R.E. Pinto, A.S. Marinho, Lipid peroxidation in mitochondrial inner membranes. I. An integrative kinetic model, Free Radical Biology and Medicine. 21 (1996) 21(7):917–943. doi:10.1016/S0891-5849(96)00185-2.

[26] F. Antunes, A. Salvador, R.E. Pinto, PHGPx and phospholipase A2/GPx: comparative importance on the reduction of hydroperoxides in rat liver mitochondria, Free Radical Biology and Medicine. 19 (1995) 669–677. doi:10.1016/0891-5849(95)00040-5.

[27] A. Salvador, J. Sousa, R.E. Pinto, Hydroperoxyl, superoxide and pH gradients in the mitochondrial matrix: a theoretical assessment, Free Radical Biology and Medicine. 31 (2001) 1208–1215.

[28] G.R. Buettner, Moving free radical and redox biology ahead in the next decade(s), Free Radical Biology and Medicine. 78 (2015) 236–238. doi:10.1016/j.freeradbiomed.2014.10.578.

[29] P.M.B.M. Coelho, A. Salvador, M.A. Savageau, Quantifying Global Tolerance of Biochemical Systems: Design Implications for Moiety-Transfer Cycles, PLoS Computational Biology. 5 (2009) e1000319. doi:10.1371/journal.pcbi.1000319.

[30] M.A. Savageau, P.M.B.M. Coelho, R.A. Fasani, D.A. Tolla, A. Salvador, Phenotypes and tolerances in the design space of biochemical systems, Proceedings of the National Academy of Sciences. 106 (2009) 6435–6440. doi:10.1073/pnas.0809869106.

[31] P.M.B.M. Coelho, A. Salvador, M.A. Savageau, Relating Mutant Genotype to Phenotype via Quantitative Behavior of the NADPH Redox Cycle in Human Erythrocytes, PLoS ONE. 5 (2010) e13031. doi:10.1371/journal.pone.0013031.

[32] A. Salvador, M.A. Savageau, Quantitative evolutionary design of glucose 6-phosphate dehydrogenase expression in human erythrocytes, Proceedings of the National Academy of Sciences. 100 (2003) 14463–14468. doi:10.1073/pnas.2335687100.

[33] A. Salvador, M.A. Savageau, Evolution of enzymes in a series is driven by dissimilar functional demands, Proceedings of the National Academy of Sciences. 103 (2006) 2226–2231. doi:10.1073/pnas.0510776103.

[34] A. Kuehne, H. Emmert, J. Soehle, M. Winnefeld, F. Fischer, H. Wenck, S. Gallinat, L. Terstegen, R. Lucius, J. Hildebrand, N. Zamboni, Acute Activation of Oxidative Pentose Phosphate Pathway as First-Line Response to Oxidative Stress in Human Skin Cells., Molecular Cell. 59 (2015) 359–371. doi:10.1016/j.molcel.2015.06.017.

[35] R.A. Poynton, A. V Peskin, A.C. Haynes, W.T. Lowther, M.B. Hampton, C.C. Winterbourn, Kinetic analysis of structural influences on the susceptibility of peroxiredoxins 2 and 3 to hyperoxidation, Biochemical Journal. 473 (2016) 411–421. doi:10.1042/BJ20150572.

[36] T.S. Chang, W. Jeong, H.A. Woo, S.M. Lee, S. Park, S.G. Rhee, Characterization of Mammalian Sulfiredoxin and Its Reactivation of Hyperoxidized Peroxiredoxin through Reduction of Cysteine Sulfinic Acid in the Active Site to Cysteine, Journal of Biological Chemistry. 279 (2004) 50994–51001. doi:10.1074/jbc.M409482200.

[37] X. Roussel, S. Boukhenouna, S. Rahuel-Clermont, G. Branlant, The rate-limiting step of sulfiredoxin is associated with the transfer of the γ-phosphate of ATP to the sulfinic acid of overoxidized typical 2-Cys peroxiredoxins, FEBS Letters. 585 (2011) 574–578. doi:10.1016/j.febslet.2011.01.012.

[38] E.S. Arnér, A. Holmgren, Physiological functions of thioredoxin and thioredoxin reductase, European Journal of Biochemistry. 267 (2000) 6102–6109. doi:10.1046/j.1432-1327.2000.01701.x.

[39] S. Urig, J. Lieske, K. Fritz-Wolf, A. Irmler, K. Becker, Truncated mutants of human thioredoxin reductase 1 do not exhibit glutathione reductase activity, FEBS Letters. 580 (2006) 3595–3600. doi:10.1016/j.febslet.2006.05.038.

[40] M. Cebula, E.E. Schmidt, E.S. Arnér, TrxR1 as a Potent Regulator of the Nrf2-Keap1 Response System, Antioxidants & Redox Signaling. 23 (2015) 823–853. doi:10.1089/ars.2015.6378.

[41] J.W. Park, J.J. Mieyal, S.G. Rhee, P.B. Chock, Deglutathionylation of 2-Cys Peroxiredoxin Is Specifically Catalyzed by Sulfiredoxin, The Journal of Biological Chemistry. 284 (2009) 23364–23374. doi:10.1074/jbc.M109.021394.

[42] A. V. Peskin, P.E. Pace, J.B. Behring, L.N. Paton, M. Soethoudt, M.M. Bachschmid, C.C. Winterbourn, Glutathionylation of the Active Site Cysteines of Peroxiredoxin 2 and Recycling by Glutaredoxin, The Journal of Biological Chemistry. 291 (2016) 3053–3062. doi:10.1074/jbc.M115.692798.

[43] Y. Du, H. Zhang, J. Lu, A. Holmgren, Glutathione and glutaredoxin act as a backup of human thioredoxin reductase 1 to reduce thioredoxin 1 preventing cell death by aurothioglucose, Journal of Biological Chemistry. 287 (2012) 38210–38219. doi:10.1074/jbc.M112.392225.

[44] W.H. Koppenol, Thermodynamics of reactions involving nitrogen-oxygen compounds, Methods in Enzymology. 268 (1996) 7–12. doi:10.1016/S0076-6879(96)68005-7.

[45] J.B. Lim, B.K. Huang, W.M. Deen, H.D. Sikes, Analysis of the lifetime and spatial localization of Hydrogen peroxide generated in the cytosol using a reduced kinetic model., Free Radical Biology and Medicine. 89 (2015) 47–53. doi:10.1016/j.freeradbiomed.2015.07.009.

[46] A. Salvador, Synergism analysis of biochemical systems. I. Conceptual framework, Mathematical Biosciences. 163 (2000) 105–129. doi:10.1016/S0025-5564(99)00056-5.

[47] A. Salvador, Synergism analysis of biochemical systems. II. Tensor formulation and treatment of stoichiometric constraints, Mathematical Biosciences. 163 (2000) 131–158. doi:10.1016/S0025-5564(99)00057-7.

[48] I. Wolfram Research, Mathematica, Version 7., Wolfram Research, Inc., 2008.

[49] X. Roussel, A. Kriznik, C. Richard, S. Rahuel-Clermont, G. Branlant, Catalytic Mechanism of Sulfiredoxin from Saccharomyces cerevisiae Passes through an Oxidized Disulfide Sulfiredoxin Intermediate That Is Reduced by Thioredoxin, Journal of Biological Chemistry. 284 (2009) 33048–33055. doi:10.1074/jbc.M109.035352.

[50] S.G. Rhee, W. Jeong, T.S. Chang, H.A. Woo, Sulfiredoxin, the cysteine sulfinic acid reductase specific to 2-Cys peroxiredoxin: its discovery, mechanism of action, and biological significance, Kidney International. 72 (2007) S3–S8. doi:10.1038/sj.ki.5002380.

[51] X. Roussel, G. Béchade, A. Kriznik, A. Van Dorsselaer, S. Sanglier-Cianferani, G. Branlant, S. Rahuel-Clermont, Evidence for the Formation of a Covalent Thiosulfinate Intermediate with Peroxiredoxin in the Catalytic Mechanism of Sulfiredoxin, Journal of Biological Chemistry. 283 (2008) 22371–22382. doi:10.1074/jbc.M800493200.

[52] J.J. Tyson, K.C. Chen, B. Novak, Sniffers, buzzers, toggles and blinkers: dynamics of regulatory and signaling pathways in the cell, Current Opinion In Cell Biology. 15 (2003) 221–231. doi:10.1016/s0955-0674(03)00017-6.

[53] T. Geiger, A. Wehner, C. Schaab, J. Cox, M. Mann, Comparative Proteomic Analysis of Eleven Common Cell Lines Reveals Ubiquitous but Varying Expression of Most Proteins, Molecular & Cellular Proteomics. 11 (2012) M111.014050–M111.014050. doi:10.1074/mcp.M111.014050.

[54] J.R. Wiśniewski, A. Vildhede, A. Norén, P. Artursson, In-depth quantitative analysis and comparison of the human hepatocyte and hepatoma cell line HepG2 proteomes, Journal of Proteomics. 136 (2016) 234–247. doi:10.1016/j.jprot.2016.01.016.

[55] R.M. Johnson, Y.-S. Ho, D.-Y. Yu, F.A. Kuypers, Y. Ravindranath, G.W. Goyette, The effects of disruption of genes for peroxiredoxin-2, glutathione peroxidase-1, and catalase on erythrocyte oxidative metabolism, Free Radical Biology and Medicine. 48 (2010) 519–525. doi:10.1016/j.freeradbiomed.2009.11.021.

[56] H.S. Jacob, S.H. Ingbar, J.H. Jandl, Oxidative hemolysis and erythrocyte metabolism in hereditary acatalasia, Journal of Clinical Investigation. 44 (1965) 1187–1199. doi:10.1172/JCI105225.

[57] C.S. Cho, G.J. Kato, S.H. Yang, S.W. Bae, J.S. Lee, M.T. Gladwin, S.G. Rhee, Hydroxyurea-Induced Expression of Glutathione Peroxidase 1 in Red Blood Cells of Individuals with Sickle Cell Anemia, Antioxidants and Redox Signaling. 13 (2010) 1–11. doi:10.1089/ars.2009.2978.

[58] G.P. Bienert, A.L.B. Møller, K.A. Kristiansen, A. Schulz, I.M. Møller, J.K. Schjoerring, T.P. Jahn, Specific aquaporins facilitate the diffusion of hydrogen peroxide across membranes., The Journal of Biological Chemistry. 282 (2007) 1183–92. doi:10.1074/jbc.M603761200.

[59] M. Jelcic, B. Enyedi, J.B. Xavier, P. Niethammer, Image-Based Measurement of H2O2 Reaction-Diffusion in Wounded Zebrafish Larvae, Biophysical Journal. 112 (2017) 2011–2018. doi:10.1016/j.bpj.2017.03.021.

[60] M. Chevallet, E. Wagner, S. Luche, A. van Dorsselaer, E. Leize-Wagner, T. Rabilloud, Regeneration of peroxiredoxins during recovery after oxidative stress - Only some overoxidized peroxiredoxins can be reduced during recovery after oxidative stress, Journal Of Biological Chemistry. 278 (2003) 37146–37153. doi:10.1074/jbc.M305161200.

[61] S.J. Montano, J. Lu, T.N. Gustafsson, A. Holmgren, Activity assays of mammalian thioredoxin and thioredoxin reductase: Fluorescent disulfide substrates, mechanisms, and use with tissue samples, Analytical Biochemistry. 449 (2014) 139–146. doi:10.1016/J.AB.2013.12.025.

[62] M. Kontou, C. Adelfalk, M. Hirsch-Kauffmann, M. Schweiger, Suboptimal Action of NF-κB in Fanconi Anemia Cells Results from Low Levels of Thioredoxin, Biological Chemistry. 384 (2003). doi:10.1515/BC.2003.166.

[63] S.J. Wei, A. Botero, K. Hirota, C.M. Bradbury, S. Markovina, A. Laszlo, D.R. Spitz, P.C. Goswami, J. Yodoi, D. Gius, Thioredoxin nuclear translocation and interaction with redox factor-1 activates the activator protein-1 transcription factor in response to ionizing radiation., Cancer Research. 60 (2000) 6688–95.

[64] P. Schroeder, R. Popp, B. Wiegand, J. Altschmied, J. Haendeler, Nuclear redox-signaling is essential for apoptosis inhibition in endothelial cells--important role for nuclear thioredoxin-1., Arteriosclerosis, Thrombosis, and Vascular Biology. 27 (2007) 2325–31. doi:10.1161/ATVBAHA.107.149419.

[65] S. Lee, S.M. Kim, R.T. Lee, Thioredoxin and Thioredoxin Target Proteins: From Molecular Mechanisms to Functional Significance, Antioxidants and Redox Signaling. 18 (2012) 1165–1207. doi:10.1089/ars.2011.4322.

[66] B.A. Wagner, J.R. Witmer, T.J. van’t Erve, G.R. Buettner, An Assay for the Rate of Removal of Extracellular Hydrogen Peroxide by Cells., Redox Biology. 1 (2013) 210–217. doi:10.1016/j.redox.2013.01.011.

[67] C.M. Doskey, V. Buranasudja, B.A. Wagner, J.G. Wilkes, J. Du, J.J. Cullen, G.R. Buettner, Tumor cells have decreased ability to metabolize H2O2: Implications for pharmacological ascorbate in cancer therapy, Redox Biology. 10 (2016) 274–284. doi:10.1016/j.redox.2016.10.010.

[68] B.K. Huang, H.D. Sikes, Quantifying intracellular hydrogen peroxide perturbations in terms of concentration, Redox Biology. 2 (2014) 955–962. doi:10.1016/j.redox.2014.08.001.

[69] F. Antunes, E. Cadenas, Cellular titration of apoptosis with steady state concentrations of H2O2: submicromolar levels of H2O2 induce apoptosis through Fenton chemistry independent of the cellular thiol state, Free Radical Biology and Medicine. 30 (2001) 1008–1018. doi:10.1016/S0891-5849(01)00493-2.

[70] F. Antunes, E. Cadenas, Estimation of H2O2 gradients across biomembranes, FEBS Letters. 475 (2000) 121–126. doi:10.1016/S0003-9861(02)00049-8.

[71] M. Saitoh, H. Nishitoh, M. Fujii, K. Takeda, K. Tobiume, Y. Sawada, M. Kawabata, K. Miyazono, H. Ichijo, Mammalian thioredoxin is a direct inhibitor of apoptosis signal-regulating kinase (ASK) 1, The EMBO Journal. 17 (1998) 2596–2606. doi:10.1093/emboj/17.9.2596.

[72] D. Kosek, S. Kylarova, K. Psenakova, L. Rezabkova, P. Herman, J. Vecer, V. Obsilova, T. Obsil, Biophysical and structural characterization of the thioredoxin-binding domain of protein kinase ASK1 and its interaction with reduced thioredoxin., The Journal of Biological Chemistry. 289 (2014) 24463–74. doi:10.1074/jbc.M114.583807.

[73] S. Kylarova, D. Kosek, O. Petrvalska, K. Psenakova, P. Man, J. Vecer, P. Herman, V. Obsilova, T. Obsil, Cysteine residues mediate high-affinity binding of thioredoxin to ASK1, The FEBS Journal. 283 (2016) 3821–3838. doi:10.1111/febs.13893.

[74] T. Obsil, V. Obsilova, Structural aspects of protein kinase ASK1 regulation, Advances in Biological Regulation. (2017) 0–1. doi:10.1016/j.jbior.2017.10.002.

[75] T. Nishida, K. Hattori, K. Watanabe, The regulatory and signaling mechanisms of the ASK family, Advances in Biological Regulation. (2017) 1–21. doi:10.1016/j.jbior.2017.05.004.

[76] L. Flohé, Changing paradigms in thiology from antioxidant defense toward redox regulation, in: Methods Enzymol, 2010: pp. 1–39. doi:10.1016/S0076-6879(10)73001-9.

[77] H.J. Forman, M. Maiorino, F. Ursini, Signaling Functions of Reactive Oxygen Species, Biochemistry. 49 (2010) 835–842. doi:10.1021/bi9020378.

[78] M.C. Sobotta, W. Liou, S. Stöcker, D. Talwar, M. Oehler, T. Ruppert, A.N. Scharf, T.P. Dick, Peroxiredoxin-2 and STAT3 form a redox relay for H2O2 signaling, Nature Chemical Biology. 11 (2014) 64–70. doi:10.1038/nchembio.1695.

[79] C. Little, P.J. O’brien, Mechanism of Peroxide-Inactivation of the Sulphydryl Enzyme Glyceraldehyde-3-Phosphate Dehydrogenase, European Journal of Biochemistry. 10 (1969) 533–538. doi:10.1111/j.1432-1033.1969.tb00721.x.

[80] C.C. Winterbourn, M.B. Hampton, Thiol chemistry and specificity in redox signaling, Free Radical Biology and Medicine. 45 (2008) 549–561. doi:10.1016/j.freeradbiomed.2008.05.004.

[81] H.S. Marinho, C. Real, L. Cyrne, H. Soares, F. Antunes, Hydrogen peroxide sensing, signaling and regulation of transcription factors, Redox Biology. 2 (2014) 535–562. doi:10.1016/j.redox.2014.02.006.

[82] D. Peralta, A.K. Bronowska, B. Morgan, É. Dóka, K. Van Laer, P. Nagy, F. Gräter, T.P. Dick, A proton relay enhances H2O2 sensitivity of GAPDH to facilitate metabolic adaptation., Nature Chemical Biology. 11 (2015) 156–63. doi:10.1038/nchembio.1720.

[83] U. Schwertassek, A. Haque, N. Krishnan, R. Greiner, L. Weingarten, T.P. Dick, N.K. Tonks, Reactivation of oxidized PTP1B and PTEN by thioredoxin 1, FEBS Journal. 281 (2014) 3545–3558. doi:10.1111/febs.12898.

[84] J.M. Denu, K.G. Tanner, Specific and Reversible Inactivation of Protein Tyrosine Phosphatases by Hydrogen Peroxide: Evidence for a Sulfenic Acid Intermediate and Implications for Redox Regulation, Biochemistry. 37 (1998) 5633–5642. doi:10.1021/bi973035t.

[85] H. Zhou, H. Singh, Z.D. Parsons, S.M. Lewis, S. Bhattacharya, D.R. Seiner, J.N. LaButti, T.J. Reilly, J.J. Tanner, K.S. Gates, The Biological Buffer Bicarbonate/CO2 Potentiates H2O2 - Mediated Inactivation of Protein Tyrosine Phosphatases, Journal of the American Chemical Society. 133 (2011) 15803–15805. doi:10.1021/ja2077137.

[86] F. Antunes, P.M. Brito, Quantitative biology of hydrogen peroxide signaling, Redox Biology. (2017). doi:10.1016/j.redox.2017.04.039.

[87] B. Medicherla, A.L. Goldberg, Heat shock and oxygen radicals stimulate ubiquitin-dependent degradation mainly of newly synthesized proteins., The Journal of Cell Biology. 182 (2008) 663–73. doi:10.1083/jcb.200803022.

[88] S. Dukan, A. Farewell, M. Ballesteros, F. Taddei, M. Radman, T. Nystrom, Protein oxidation in response to increased transcriptional or translational errors, Proceedings Of The National Academy Of Sciences Of The United States Of America. 97 (2000) 5746–5749. doi:10.1073/pnas.100422497.

[89] H.H. Jang, K.O. Lee, Y.H. Chi, B.G. Jung, S.K. Park, J.H. Park, J.R. Lee, S.S. Lee, J.C. Moon, J.W. Yun, Y.O. Choi, W.Y. Kim, J.S. Kang, G.-W. Cheong, D.-J. Yun, S.G. Rhee, M.J. Cho, S.Y. Lee, Two enzymes in one. Two yeast peroxiredoxins display oxidative stress-dependent switching from a peroxidase to a molecular chaperone function., Cell. 117 (2004) 625–35. doi:10.1016/j.cell.2004.05.002.

[90] J.C. Moon, Y.-S. Hah, W.Y. Kim, B.G. Jung, H.H. Jang, J.R. Lee, S.Y. Kim, Y.M. Lee, M.G. Jeon, C.W. Kim, M.J. Cho, S.Y. Lee, Oxidative stress-dependent structural and functional switching of a human 2-Cys peroxiredoxin isotype II that enhances HeLa cell resistance to H2O2-induced cell death, Journal of Biological Chemistry. 280 (2005) 28775–28784. doi:10.1074/jbc.M505362200.

[91] J.C. Lim, H.-I. Choi, Y.S. Park, H.W. Nam, H.A. Woo, K.-S. Kwon, Y.S. Kim, S.G. Rhee, K. Kim, H.Z. Chae, Irreversible oxidation of the active-site cysteine of peroxiredoxin to cysteine sulfonic acid for enhanced molecular chaperone activity., The Journal of Biological Chemistry. 283 (2008) 28873–80. doi:10.1074/jbc.M804087200.

[92] J.H. Hoffmann, K. Linke, P.C.F. Graf, H. Lilie, U. Jakob, Identification of a redox-regulated chaperone network, The EMBO Journal. 23 (2003) 160 LP-168. doi:10.1038/sj.emboj.7600016.

[93] S. Hanzén, K. Vielfort, J. Yang, F. Roger, V. Andersson, S. Zamarbide-Forés, R. Andersson, L. Malm, G. Palais, B. Biteau, B. Liu, M.B.B. Toledano, M. Molin, T. Nyström, Lifespan Control by Redox-Dependent Recruitment of Chaperones to Misfolded Proteins, Cell. 166 (2016) 140–151. doi:10.1016/j.cell.2016.05.006.

[94] T.J. Phalen, K. Weirather, P.B. Deming, V. Anathy, A.K. Howe, A. van der Vliet, T.J. Jönsson, L.B. Poole, N.H. Heintz, Oxidation state governs structural transitions in peroxiredoxin II that correlate with cell cycle arrest and recovery, Journal of Cell Biology. 175 (2006) 779–789. doi:10.1083/jcb.200606005.

[95] A.M. Day, J.D. Brown, S.R. Taylor, J.D. Rand, B.A. Morgan, E.A. Veal, Inactivation of a Peroxiredoxin by Hydrogen Peroxide Is Critical for Thioredoxin-Mediated Repair of Oxidized Proteins and Cell Survival, Molecular Cell. 45 (2012) 398–408. doi:10.1016/j.molcel.2011.11.027.

[96] S. Salzano, P. Checconi, E.-M. Hanschmann, C.H. Lillig, L.D. Bowler, P. Chan, D. Vaudry, M. Mengozzi, L. Coppo, S. Sacre, K.R. Atkuri, B. Sahaf, L.A. Herzenberg, L.A. Herzenberg, L. Mullen, P. Ghezzi, Linkage of inflammation and oxidative stress via release of glutathionylated peroxiredoxin-2, which acts as a danger signal., Proceedings of the National Academy of Sciences of the United States of America. 111 (2014) 12157–62. doi:10.1073/pnas.1401712111.

[97] L. Mullen, E.-M. Hanschmann, C.H. Lillig, L.A. Herzenberg, P. Ghezzi, Cysteine Oxidation Targets Peroxiredoxins 1 and 2 for Exosomal Release through a Novel Mechanism of Redox-Dependent Secretion., Molecular Medicine (Cambridge, Mass.). 21 (2015) 98–108. doi:10.2119/molmed.2015.00033.

[98] P. Checconi, S. Salzano, L. Bowler, L. Mullen, M. Mengozzi, E.-M. Hanschmann, C.H. Lillig, R. Sgarbanti, S. Panella, L. Nencioni, A.T. Palamara, P. Ghezzi, Redox Proteomics of the Inflammatory Secretome Identifies a Common Set of Redoxins and Other Glutathionylated Proteins Released in Inflammation, Influenza Virus Infection and Oxidative Stress, PLOS ONE. 10 (2015) e0127086. doi:10.1371/journal.pone.0127086.

[99] M. Mishra, H. Jiang, L. Wu, H.A. Chawsheen, Q. Wei, The sulfiredoxin-peroxiredoxin (Srx-Prx) axis in cell signal transduction and cancer development., Cancer Letters. 366 (2015) 150–9. doi:10.1016/j.canlet.2015.07.002.

[100] A. Mukherjee, S.G. Martin, The thioredoxin system: a key target in tumour and endothelial cells, The British Journal of Radiology. 81 (2008) S57–S68. doi:10.1259/bjr/34180435.

[101] R. Bretón-Romero, S. Lamas, Hydrogen peroxide signaling in vascular endothelial cells., Redox Biology. 2 (2014) 529–534. doi:10.1016/j.redox.2014.02.005.

[102] C.A. Tairum Jr., M.A. de Oliveira, B.B. Horta, F.J. Zara, L.E.S. Netto, Disulfide Biochemistry in 2-Cys Peroxiredoxin: Requirement of Glu50 and Arg146 for the Reduction of Yeast Tsa1 by Thioredoxin, Journal of Molecular Biology. 424 (2012) 28–41. doi:10.1016/j.jmb.2012.09.008.

[103] A. V Peskin, N. Dickerhof, R.A. Poynton, L.N. Paton, P.E. Pace, M.B. Hampton, C.C. Winterbourn, Hyperoxidation of peroxiredoxins 2 and 3: Rate constants for the reactions of the sulfenic acid of the peroxidatic cysteine, The Journal of Biological Chemistry. 288 (2013) 14170–14177. doi:10.1074/jbc.M113.460881.

